# A Protein Disulfide Isomerase Coordinates Redox Homeostasis and ER Calcium Regulation for Optimal Lytic Cycle Progression in *Toxoplasma gondii*

**DOI:** 10.1101/2025.05.15.654346

**Authors:** Katherine E Moen, Silvia N J Moreno

## Abstract

The endoplasmic reticulum (ER) maintains an oxidative environment that facilitates disulfide bond formation, a critical process for proper protein folding. Protein disulfide isomerases (PDIs) are ER resident enzymes that facilitate the formation, breakage, and rearrangement of disulfide bonds between cysteine residues, thereby stabilizing protein structures. Although PDIs are functionally diverse, they all contain at least 1 thioredoxin-like domain and mediate disulfide exchange through their conserved CXXC motifs. The Apicomplexan parasite, *Toxoplasma gondii*, infects approximately one third of the world population, posing a significant risk to immunosuppressed individuals and unborn fetuses. The fast-replicating tachyzoite form engages in a lytic cycle, causing host tissue damage and contributing to pathogenesis. While approximately 26 PDIs are predicted to be present in *T. gondii*, their specific roles remain largely unexplored. In this study, we investigate TgPDIA3, a *T. gondii* PDI localized to the ER, along with several of its interacting protein substrates. We explore its role in ER redox activity and calcium sequestration and assess how these functions contribute to the parasite’s lytic cycle.

**Importance:** The lytic cycle of *Toxoplasma gondii* is essential to the pathogenesis of toxoplasmosis, with calcium signaling playing a crucial role in driving this process. Cytosolic calcium is tightly regulated through either sequestration into intracellular stores or extrusion from the cell. The ER, likely the largest calcium store in *T. gondii*, remains poorly characterized. In this study, we identify a link between ER redox regulation and calcium homeostasis and signaling in *T. gondii*. These findings suggest that redox-controlled calcium homeostasis and flux in the ER is a key driver of the parasite’s lytic cycle progression.

## Introduction

*Toxoplasma gondii* is an obligate intracellular Apicomplexan parasite that infects approximately one-third of the world’s human population [2]. In the US there is an estimated seroprevalence of 11% within individuals 6 years and older [3]. Reactivation of *T. gondii* latent infections in individuals who become immunocompromised can cause toxoplasmic encephalitis [4, 5]. Congenital transmission can occur when women become newly infected during pregnancy [6]. During the acute infection, the fast-growing tachyzoite engages in a lytic cycle, during which it attaches and actively invades host cells, undergoes asexual replication inside a parasitophorous vacuole (PV), followed by egress, rupturing the host cell’s membranes. This lytic cycle drives the parasite’s pathogenesis by continuously disrupting host tissues and spreading through all host organs [7, 8]. *T. gondii* possess specialized organelles called micronemes and rhoptries, unique to apicomplexans, that secrete microneme and rhoptry proteins, respectively. Microneme proteins are secreted during egress, gliding, host cell attachment, and invasion [9]. After attachment rhoptry neck (RON) and rhoptry bulb (ROP) proteins are secreted to facilitate parasite invasion into the host cell and to participate in the development of the PV [10]. Increasing cytosolic calcium using ionophores triggers microneme secretion, a crucial event for most transitions in the parasites’ lytic cycle [11].

Calcium signaling is universal and affects all eukaryotic cells [12, 13]. However, cells must maintain low resting level of cytosolic calcium (<100 nM) because it can form toxic precipitates with phosphates in the cytosol. Cells sequester calcium in intracellular stores, particularly the endoplasmic reticulum (ER) which serves as the largest intracellular calcium store in most eukaryotic cells [13]. ER calcium homeostasis is maintained by calcium-binding proteins (CBPs) within the ER lumen, in addition to calcium channels, pumps, and exchangers located on the ER membrane. The sarco-endoplasmic reticulum calcium ATPase (SERCA) pumps calcium into the ER and helps maintaining low cytosolic levels of calcium. In mammals, the inositol trisphosphate receptor (IP_3_R) and the ryanodine receptor (RyR) are channels embedded in the ER membrane that, when activated, release calcium into the cytosol to initiate downstream signaling [12].

The high concentration of calcium in the ER is essential for its multiple functions, including signaling, protein chaperoning, and maintaining redox homeostasis. The ER serves as the cell’s quality control center for post-translational processing of membrane proteins [14] and proteins destined for secretion [15]. Molecular chaperones use hydrophobic interactions and ATP to fold nascent polypeptides into their correct three-dimensional conformations [16]. Binding immunoglobin protein (BiP), an ER resident HSP70 molecular chaperone, binds calcium for sensing and storage, and its chaperone activity is regulated by calcium binding [17, 18]. Lectin chaperones in the ER play a crucial role in folding glycoproteins and often function as low-affinity, high-capacity CBPs, essential for calcium sensing and storage. Calcium binding also modulates their chaperone functions [18].

The ER is an oxidizing environment [19] and, as such, serves as a major site of oxidative protein folding within the cell. This post-translational process, carried out by ER redox active enzymes consists of the formation, cleavage, and rearrangement of disulfide bonds in client proteins, stabilizing them in their correct conformations. Protein disulfide isomerases (PDIs) are ER-resident redox enzymes that belong to the thioredoxin (TRX) superfamily of dithiol/disulfide oxidoreductases [20–23]. Redox active PDIs contain canonical CXXC motifs, which cycle between oxidized and reduced states to catalyze the formation, cleavage, or rearrangement of disulfide bonds in their protein substrates (**Fig 1A**). Humans possess 21 known PDIs, many of which are multifunctional. Most PDIs feature specialized b or b’ domains rich in hydrophobic residues that are important for substrate recognition and assist in non-redox protein folding [24–27]. Some PDIs contain low-affinity, high-capacity calcium-binding domains that contribute to ER calcium buffering and sensing [28]. Additionally, certain PDIs regulate other proteins through disulfide bond exchange. In animals, several PDIs play a role in calcium homeostasis by adding or removing regulatory disulfide bonds on SERCA or IP_3_R [28–31].

**Fig 1.**
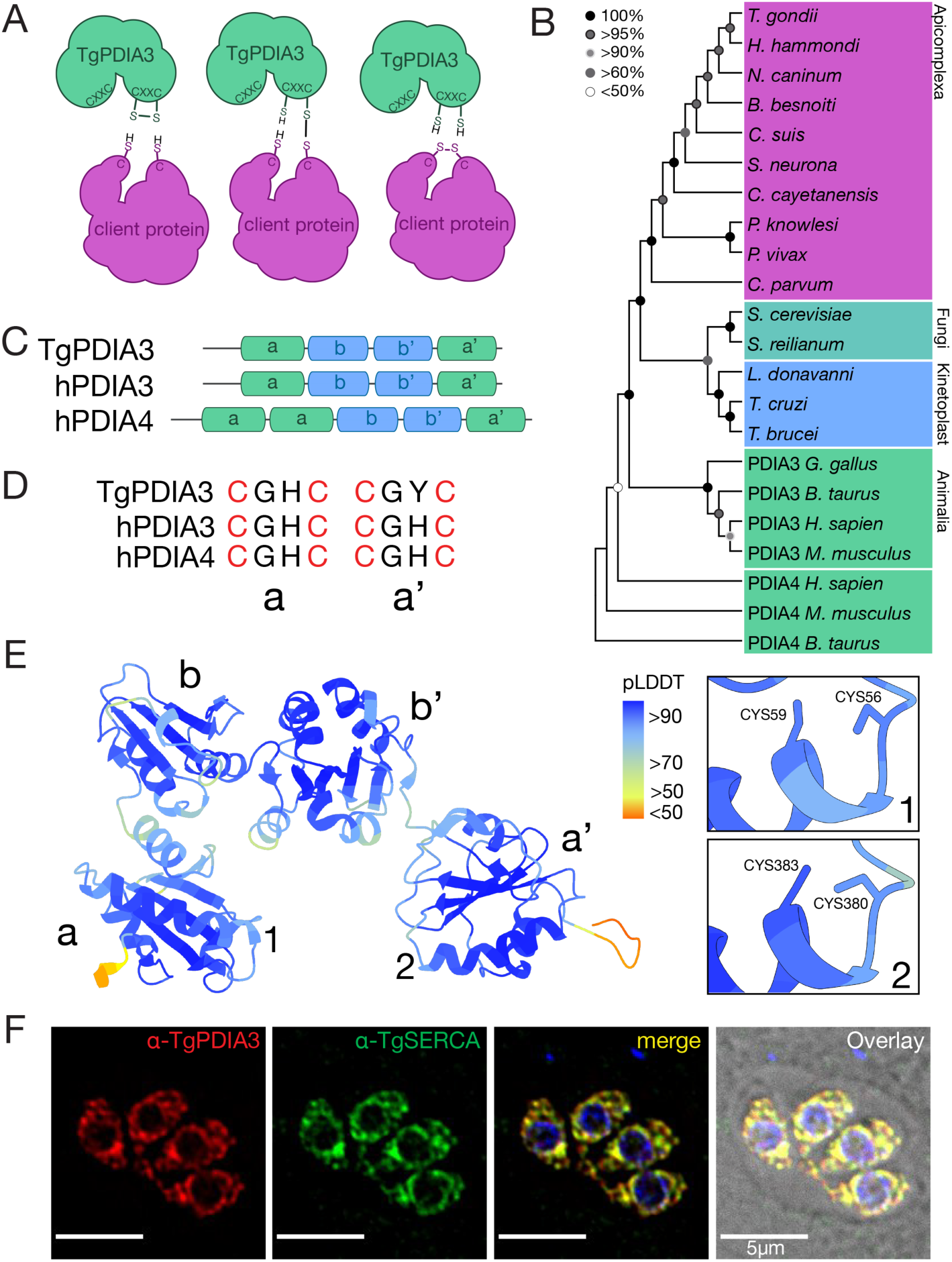
TgPDIA3 is an ER-resident PDI. A) Scheme showing the redox activity of TgPDIA3 on a client protein. B) Phylogenetic analysis of TgPDIA3 and blasted predicted orthologs, including both mammalian PDIA3 and PDIA4. The sequences used are shown in **Table S1**. C) Organization of the PDI conserved **a**, **b**, **b’** and **a’** domains in TgPDIA3, HsPDIA3, and HsPDIA4. D) CXXC motifs in the **a** and **a’** domains of TgPDIA3, HsPDIA3, and HsPDIA4. E) AlphaFold v2.0 model of TgPDIA3 indicating the two CXXC active sites (1 and 2), and the 4 globular domains with average pLDDT scores for the domains being **a**:92.42 **b**:93.54 **b’**:94.15 **a’**:94.72. F) IFA of intracellular parasites utilizing the αTgPDIA3 antibody and co-localization with αTgSERCA (scale bar: 5 μm).

In the ER, chaperoning, redox, and calcium homeostasis are interconnected and delicately balanced. ER stress can be triggered by a depletion of ER calcium, which lowers the folding capacity of the ER and leads to an accumulation of mis-folded proteins, disrupting ER redox homeostasis. Furthermore, accumulation of mis-folded proteins may lead to disruption of calcium and redox homeostasis, triggering ER stress [32, 33]. Disturbances in ER redox homeostasis can also lead to accumulation of misfolded proteins and cause ER stress [34, 35]. Some redox proteins, in collaboration with other chaperones, interact with IP_3_R at mitochondrial associated membranes (MAMs), which regulates calcium transfer to the mitochondria [29, 36, 37]. In this way, ER stress caused by excessive calcium buildup can lead to either an increase in ATP production or the initiation of apoptosis [38].

PDIs operate at the intersection of these finely balanced processes, serving as communication links among various ER functions. Nearly all PDIs are redox-active and play a pivotal role in maintaining ER redox homeostasis. While some PDIs modulate ER calcium levels through their interactions with ER calcium pumps and channels, others directly bind, sense, and sequester calcium within the ER. Beyond stabilizing client proteins with disulfide bond formation, some PDIs interact with molecular and lectin chaperones, influencing and being influenced by ER chaperone activity [27, 39, 40].

The *T. gondii* genome predicts the presence of approximately 26 PDIs. In this study, we characterized a *T. gondii* PDI, predicted to be orthologous to human PDIA3 (also known as ERp57). In animals, PDIA3 is shown to regulate SERCA activity through redox modifications [31], and is actively secreted [41]. We investigated the *T. gondii* PDI’s enzymatic activity and its involvement in various ER functions, including the maintenance of ER calcium homeostasis, interactions with potential substrates, and association with secreted effectors. We present a comprehensive functional characterization of a PDI and its substrates, highlighting their biochemical roles and consequent impact on the *T. gondii* lytic cycle.

## Results

### TgPDIA3 is an essential ER resident protein

A previous proteomic analysis of interacting partners of an endoplasmic reticulum-localized CBP identified two protein disulfide isomerases (PDIs), which are the focus of this study. We characterized *TGGT1_211680* and *TGGT1_249270*, which are predicted to encode a PDI and a putative PDI, respectively. CRISPR fitness scores for both genes as reported by Sidik et al [42] suggest that they are essential for parasite viability. We termed these proteins TgPDIA3 (TGGT1_211680) and TgPDIA6 (TGGT1_249270) with their corresponding genes named *Tgpdia3* and *Tgpdia6*. Phylogenetic analysis of TgPDIA3 (**Fig 1B and Table S1**), including BLAST comparisons with predicted orthologous genes across different kingdoms, revealed a close evolutionary relationship with human PDIA3 (aka ERp57) [41] and PDIA4 sharing approximately 34% sequence identity with each. The domain architecture of TgPDIA3 mirrors that of human PDIA3, both featuring four globular domains, while PDIA4 contains five (**Fig 1C**). The **a** catalytic domains of all 3 PDIs share the canonical CGHC active site [43]. However, the **a’** domain of TgPDIA3 displays a substitution of histidine (H) for tyrosine (Y), replacing a charged with an uncharged residue which could potentially alter its redox function (**Fig 1D**). Using AlphaFold predicted models [44, 45], we visualized the spatial orientation of TgPDIA3 catalytic motifs and found that both face inward towards each other, similar to PDIA3 [46, 47], suggesting potential functional similarities (**Figs 1E and S1A**). For TgPDIA6, phylogenetic analysis indicated the closest relationship to human PDIA6 (**Fig S2A and Table S1**), with which it shares approximately 42% sequence identity. Structural overlays of the AlphaFold models revealed high structural similarity between TgPDIA6 and PDIA6, though notable differences in domain orientations were seen (**Fig S2B-C**).

Like many ER proteins, both TgPDIA3 and TgPDIA6 possess an N-terminal signal peptide and a C-terminal retention signal, with the most common retention signals being KDEL and HDEL. Notably, TgPDIA3 features an uncommon retention signal, GEEL. Because of the presence of the retention signal, we predicted that adding a C-terminal tag could alter the localization of the protein, while an N-terminal tag could be cleaved off during maturation. Hence, we generated αTgPDIA3 antibodies in mice to visualize endogenous TgPDIA3. The *Tgpdia3* gene was first cloned into a bacterial expressing vector followed by expression and purification of the TgPDIA3 protein. This purified protein was used as an antigen for immunizing mice and generating αTgPDIA3 antibodies (**Fig S3A-B**). The specificity of these antibodies was validated by western blots of parasite lysates (**Fig S3C**). Immunofluorescence assays (IFAs) using the generated antibodies successfully visualized endogenous TgPDIA3, which localized to the ER, as confirmed by its co-localization with αTgSERCA [48], an established ER marker (**Figs 1F and S3D**). To visualize TgPDIA6, we introduced a 3-copy hemagglutinin (3xHA) epitope upstream of its endogenous KDEL retention signal. ER localization of the tagged TgPDIA6 was demonstrated through co-localization with αTgSERCA (**Fig S2D**).

With the aim to investigate the function of TgPDIA3 and TgPDIA6 in *T. gondii*, we generated conditional knockdown mutants (*iΔTgPDIA3-3Ty*, *iΔTgPDIA3* [no tag], and *iΔTgPDIA6-3Ty*) by inserting a tetracycline response element [49] upstream of a SAG4 promoter at the 5’ end of each gene (**Fig S1B**). In this system, gene expression is downregulated by adding anhydrotetracycline (ATc) to the culture media. Successful gene tagging was confirmed by PCR (**Figs S1C and S2E**) and down regulation of protein expression was validated by western blot (**Figs S1D and S2F**). To evaluate the necessity of TgPDIA3 and TgPDIA6 in the *T. gondii* lytic cycle we performed plaque assays, in which the parasite engages in cycles of invasion, replication, and egress causing host cell lysis. The resulting plaques appear as cleared zones in the host cell monolayer stained with crystal violet. Parasites lacking TgPDIA3 formed significantly smaller plaques, indicating that TgPDIA3 is crucial for at least one step of the lytic cycle (**Fig S1E-F**). Despite the CRISPR fitness score of -4.56 predicted for TgPDIA6, which suggested essentiality, the *iΔTgPDIA6-3Ty* mutant exhibited no significant differences in plaque size compared to controls (**Fig S2G-H**). To confirm these findings, we transfected both *iΔTgPDIA3* and *iΔTgPDIA6-3Ty* mutants with a cytosolic red fluorescent protein (RFP) and isolated *iΔTgPDIA3-RFP* and *iΔTgPDIA6-RFP* mutants. Growth assays measured red fluorescence over 8 days, and using a standard curve, we calculated parasite numbers based on fluorescence intensity. In the *iΔTgPDIA3- RFP* mutant, parasite growth was significantly decreased after only 4 days of ATc treatment (**Fig S1G**), supporting the essential role of TgPDIA3 for the lytic cycle. However, the *iΔTgPDIA6-RFP* mutant showed no significant growth differences compared to controls (**Fig S2I**). Further exploring the TgPDIA3 role in the lytic cycle, we performed replication assays by allowing control and *iΔTgPDIA3-RFP* parasites to invade hTERT cells. Twenty hours post infection (hpi), parasitophorous vacuoles containing 1, 2, 4, or 8+ parasites were enumerated. The *iΔTgPDIA3- RFP* mutant cultured with ATc exhibited a significant reduction in the number of PVs with 4 parasites and an increase in the number of PVs with 2 parasites compared to controls (-ATc or *TatiΔku80*), indicating that TgPDIA3 plays a significant role in parasite replication (**Fig S1H**). To investigate the role of TgPDIA3 in host cell invasion we conducted a modified red-green assay [50, 51]. After allowing invasion, extracellular RFP-expressing parasites were probed with αSAG1 to distinguish them from the invaded parasites (**Fig S1I**). The *iΔTgPDIA3-RFP* mutant pre-incubated with ATc for 48 hrs exhibited significantly reduced invaded parasites compared to controls (**Fig S1J**), indicating that TgPDIA3 is involved in the process of host cell invasion by *T. gondii* tachyzoites.

#### TgPDIA3 and TgPDIA6 catalyze the re-folding of denatured GFP in vitro

Protein folding in the ER, facilitated by molecular chaperones, is critical for achieving correct three-dimensional protein structures. Disulfide bond formation serves as a stabilizing mechanism by covalently linking cysteine residues, thereby restricting conformational flexibility and maintaining proper protein shape, essential for biological function [52].

To evaluate the ability of TgPDIA3 and TgPDIA6 to facilitate protein folding *in vitro* we utilized a recombinant GFP denatured by acid treatment as a substitute client [53]. We cloned and created recombinant soluble protein for TgPDIA3, TgPDIA6, and GFP (**Fig S4**). We acid-denatured the recombinant GFP, then exposed it to a renaturing buffer in either the absence or presence of recombinant PDI. We measured the protein’s refolding as a recovery of GFP fluorescence (**Fig 2A**). The presence of TgPDIA3 significantly enhanced the rate of GFP refolding compared to conditions lacking TgPDIA3 (**Fig 2B-C**), although it did not increase the total fluorescence gain (**Fig 2D**). TgPDIA6 was less efficient at refolding GFP, but the total fluorescence gained at lower concentrations of TgPDIA6 did increase significantly compared to the PDI free condition (**Fig 2B, 2E, and 2F**). Additionally, both TgPDIA3 and TgPDIA6 had a protective effect on the GFP as the fluorescence loss observed after the initial increase was mitigated by the presence of TgPDIA3 and even more so by the presence of TgPDIA6 (**Fig 2G-H**). In summary, TgPDIA3 significantly increased the rate of GFP refolding. While TgPDIA6 did not dramatically increase the refolding rate, the total fluorescence increase of GFP was significantly higher than the no-PDI control at lower concentrations of TgPDIA6.

**Fig 2.**
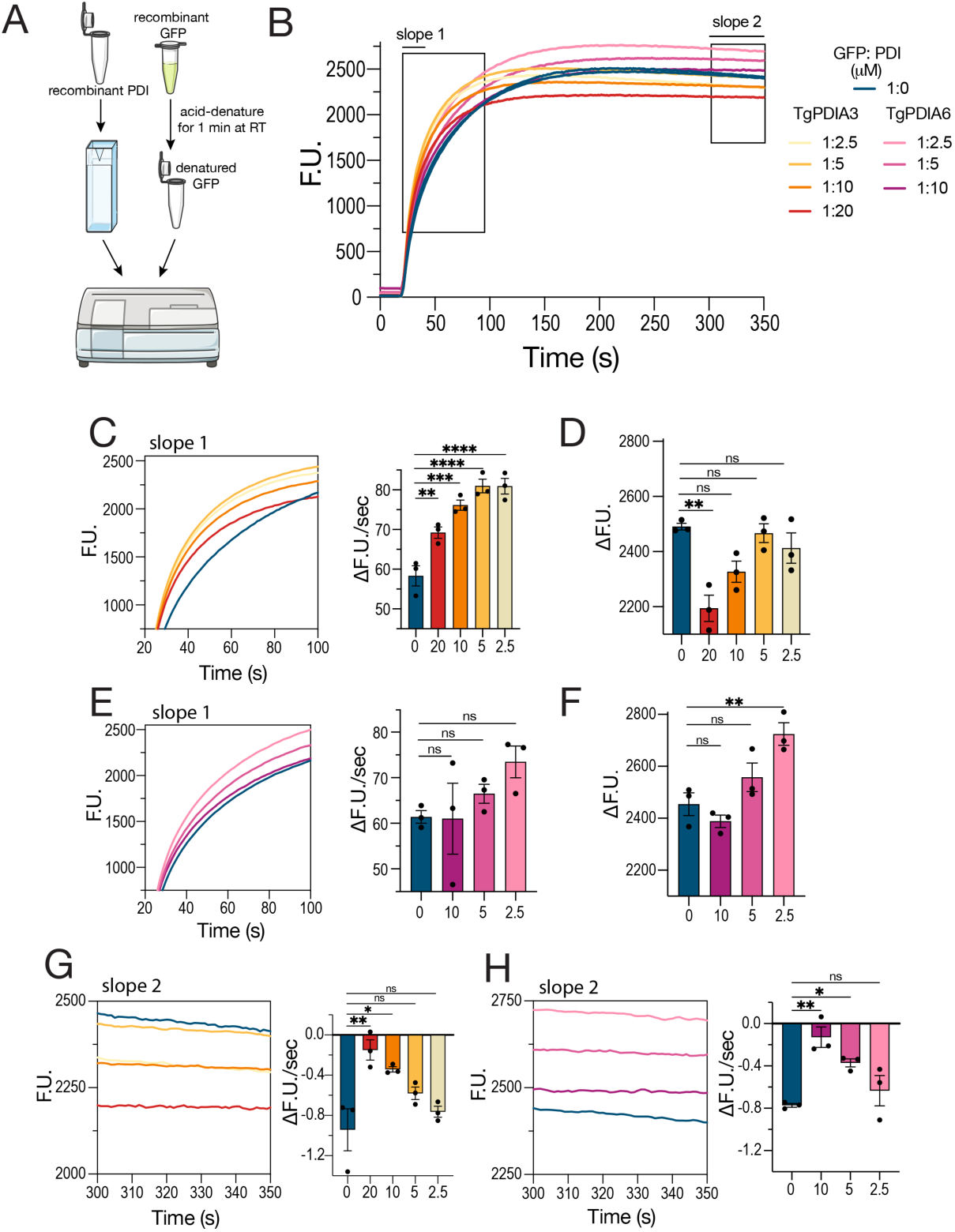
TgPDIA3 and TgPDIA6 facilitate the refolding of denatured GFP *in vitro*. A) Scheme of the protein refolding assay utilizing recombinant acid-denatured GFP and recombinant TgPDIA3 or TgPDIA6. B) Average traces showing GFP fluorescence recovery without added PDI or in the presence of different concentrations of TgPDIA3 or TgPDIA6. The numbers to the right indicate ratios GFP:PDI. C) Quantification of the initial slope of GFP fluorescence recovery in the absence or presence of varying concentrations of TgPDIA3. D) Quantification of total change in GFP fluorescence in the absence or presence of varying concentrations of TgPDIA3. E) Quantification of the initial slope of GFP fluorescence recovery in the absence or presence of varying concentrations of TgPDIA6. F) Quantification of total change in GFP fluorescence in the absence or presence of varying concentrations of TgPDIA6. G-H) Quantification of the slope of GFP fluorescence loss after initial recovery in the absence or presence of varying concentrations of TgPDIA3 (G) or TgPDIA6 (H). All quantifications using one-way ANOVA for statistical analysis (n=3).

#### TgPDIA3 is a redox-active PDI with a broad range of substrates

As a part of their primary functions, PDIs facilitate the redox folding of their client proteins. To investigate the redox role of TgPDIA3, we utilized the electrophilic crosslinker divinyl sulfone (DVSF), which covalently links cysteine residues between PDIs and their substrates during disulfide exchange, effectively trapping them together (**Fig 3A**). We incubated *TatiΔku80* parasites with or without DVSF following pre-incubation with or without N-ethylmalemide (NEM), a thiol-reactive compound that blocks disulfide bond formation and serves as a control for DVSF specificity toward cysteine residues [54–56] (**Fig 3A**).

**Fig 3.**
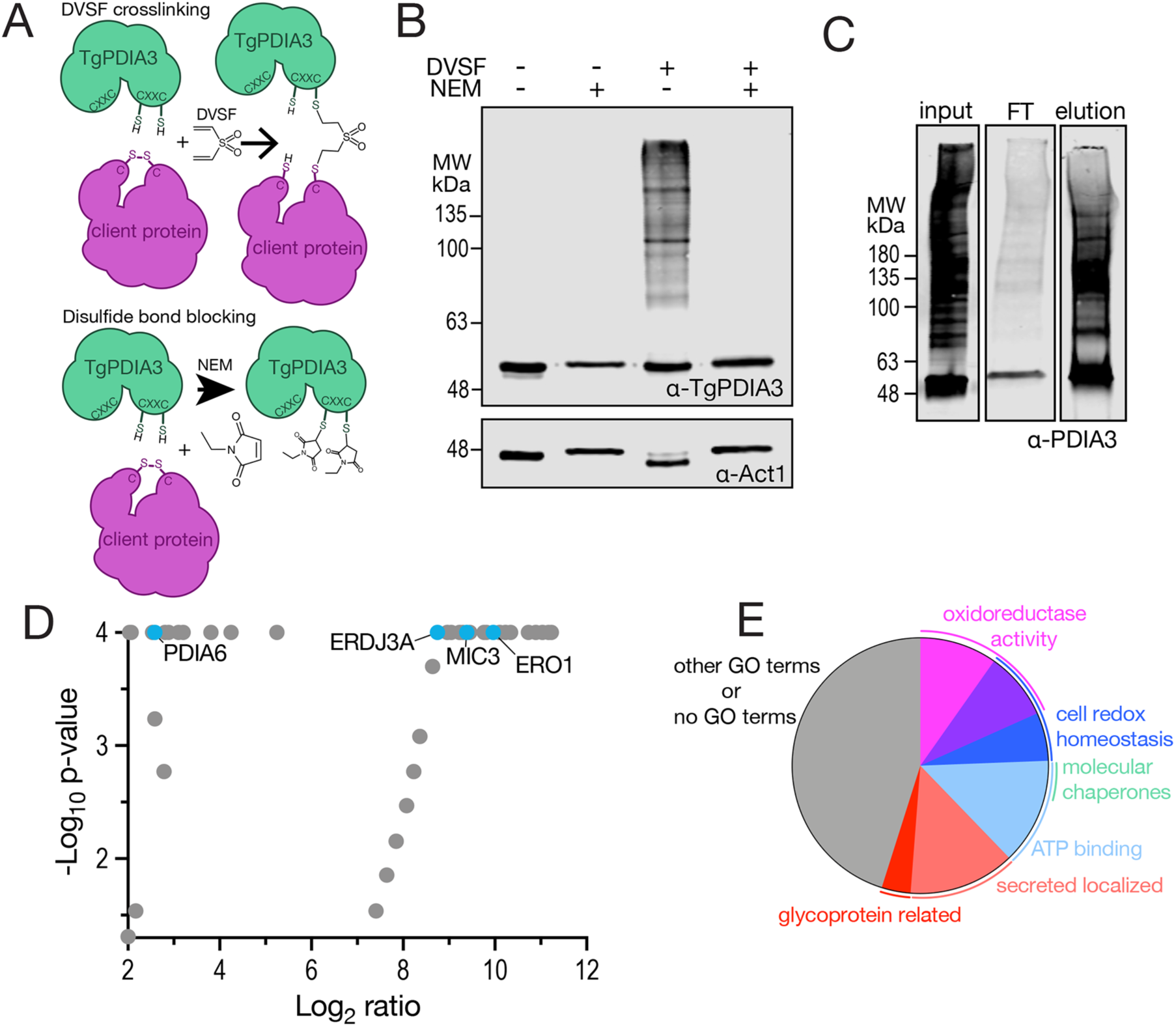
TgPDIA3 is a redox active PDI with a broad range of substrates. A) Scheme showing how DVSF covalently crosslinks PDIs to their client proteins and how pre-incubation with NEM blocks this interaction. B) Western blot analysis of parasite lysates incubated with DVSF with and without NEM, as indicated in the figure showing *iΔTgPDIA3* bound to potential client proteins (+DVSF/-NEM). ACT1 (α-actin 1) was used as loading control. C) Western blot analysis of the pre-IP input, flowthrough, and post-IP elution (all +DVSF) from the αTgPDIA3 immunoprecipiation, probed with αTgPDIA3 antibody. D) Volcano plot depicting DVSF-enriched proteins with the Log_2_ fold change (FC) (x-axis) and -Log_10_ p-value (y-axis) Fisher’s exact test was used for statistical analysis (n=3). E) pie chart of DVSF-enriched proteins grouped by LOPIT [1] predicted localization, predicted GO terms from ToxoDB, and product description.

We next analyzed the cell lysates by western blot, probing with the αTgPDIA3 antibody. In the +DVSF/-NEM condition, the observed smear suggested multiple protein interactions, which were absent under +NEM conditions, confirming the specific interaction of TgPDIA3 with a broad range of substrates (**Fig 3B**). To identify these substrates, we first incubated *TatiΔku80* parasites with DVSF, then performed immunoprecipitation of lysates using αTgPDIA3-coupled beads (**Fig 3C**), followed by liquid chromatography tandem-mass spectrometry (LC-MS/MS) analysis of the captured proteins. The *TatiΔku80* parental line, incubated without DVSF, was subjected to the same procedure and served as a negative control.

LC-MS/MS analysis revealed 82 proteins significantly enriched in the +DVSF samples compared to the -DVSF control (**Fig 3D**). Among these, 16 proteins had gene ontology (GO) terms associated with predicted oxidoreductase activity (**Fig 3E**). This group included peroxiredoxins, dehydrogenases, and ER oxidoreductin 1 (ERO1) (TGGT1_300380) an ER redox enzyme that, in mammals, interacts with PDIs (such as ERp57), to oxidize their CXXC motifs and sustain catalytic function [57] (**Fig 3D and Table 1**). Several other PDIs were also significantly enriched, including TgPDIA6, which aligns with previous findings that PDIs interact with each other [58, 59]. Interestingly, MIC3 was enriched in the +DVSF condition, suggesting that TgPDIA3 may contribute to the maturation of MIC3 through disulfide bond formation (**Fig 3D** and **Table 1**).

**Table 1:** Top 20 DVSF-enriched proteins with αTgPDIA3-IP. The 20 most enriched proteins identified based on the Log2 fold change (FC) in +DVSF compared to -DVSF. The table includes gene IDs, predicted gene products, Log2 (FC) values, and LOPIT-predicted localizations. All p-values were <0.0001 (n=3). The complete list of peptides is part of the supplemental Proteomic Results: anti TgPDIA3 IP1 and IP2.

|  | GeneID | Annotated Name | Log2FC | LOPIT |
| --- | --- | --- | --- | --- |
| 1 | TGGT1_278830 | glucose-6-phosphate 1-dehydrogenase | 11.23 | nucleus |
| 2 | TGGT1_225850 | M28 family peptidase | 11.17 | ER |
| 3 | TGGT1_227948 | M16 family peptidase (TLN2) | 11.04 | nucleolus |
| 4 | TGGT1_215910 | hypothetical protein | 10.89 | DG |
| 5 | TGGT1_228170 | IMC2A | 10.73 | DG |
| 6 | TGGT1_277270 | NTPase II | 10.34 | DG |
| 7 | TGGT1_300350 | cysteine desulfurase/selenocysteine lyase fam PLP dependent transferase | 10.23 | ER |
| 8 | TGGT1_225050 | putative adenosylhomocysteinase | 10.23 | cytosol |
| 9 | TGGT1_239752 | hypothetical protein | 10.23 | apicoplast |
| 10 | TGGT1_311210 | hypothetical protein | 10.10 | ER |
| 11 | TGGT1_271760 | seryl-tRNA synthetase (SerRS2) | 10.10 | ER |
| 12 | TGGT1_272660 | hypothetical protein | 10.10 | ER |
| 13 | TGGT1_300380 | putative ERO1 | 9.97 | ER |
| 14 | TGGT1_254490 | Sel1 repeat-containing protein | 9.81 | ER |
| 15 | TGGT1_247350 | trx domain-containing protein | 9.76 | ER |
| 16 | TGGT1_232250 | catalase | 9.45 | cytosol |
| 17 | TGGT1_247550 | HSP60 | 9.39 | Mito |
| 18 | TGGT1_319560 | MIC3 | 9.39 | MIC |
| 19 | TGGT1_270220 | hypothetical protein | 9.39 | ER |
| 20 | TGGT1_232350 | LDH1 | 9.30 | cytosol |

We next characterized two potential clients of TgPDIA3, TgERDJ3A (TGGT1_209950), a putative thioredoxin predicted to be a PDI, and ERO1, both of which were enriched in the +DVSF condition. We C-terminally HA-tagged TgERDJ3A inserting its endogenous KDEL retention signal downstream to the tag. Although LOPIT data predicted TgERDJ3A to localize to the apicoplast, it instead localized to the ER as confirmed by co-localization with αTgSERCA (**Fig S5A**). While TgERDJ3A was predicted to be essential based on its fitness score (-5.2), TgERO1 was not (fitness score 0.82). To assess whether TgERDJ3A and TgERO1 were fitness-conferring, we inserted a regulatable promoter and an N-terminal Ty-tag into each gene. The respective conditional mutants (*iΔTgERDJ3A* and *i11TgERO1)* were generated, and their growth phenotypes were analyzed using plaque assays. TgERDJ3A was found to play a significant role in parasite growth, as evidenced by the markedly smaller plaques formed by the *iΔERDJ3A* mutant upon addition of ATc to the culture media. In contrast TgERO1 was confirmed to be non-essential as the *i11ERO1* mutant was unaffected by the presence of ATc (**Fig S5B-D**). While TgERO1 was predicted to contain transmembrane domains, TgPDIA6 and TgERDJ3A were predicted to be soluble. We performed membrane extractions to investigate further these predictions (**Fig S5E**). We incubated the tagged mutants (*TgPDI6-3Ty*, *TgERDJ3A-3Ty* and *TgERO1-3Ty*) with DVSF and analyzed membrane and soluble fractions of each lysate by western blots. The results revealed a unique banding pattern of substrates for each mutant, indicating distinct binding partners and suggesting that specific PDIs may have unique redox regulatory roles in modulating their client proteins (**Fig S5E**).

To further characterize the TgPDIA3 interactome beyond its redox substrates, we C-terminally tagged TgPDIA3 with a modified TurboID plasmid. To minimize potential mislocalization we inserted the protein’s endogenous GEEL retention signal downstream to the TurboID sequence (**Fig S6A**). The TgPDIA3-TID clonal mutant was successfully isolated, and the tagging was validated through western blot and IFAs with αHA antibody (**Fig S6B-C**).

To confirm the functionality of the TurboID, parasites were incubated with biotin for 30, 60, and 90 minutes, followed by western blot analysis with streptavidin. We observed a time- increase in biotinylation in the TgPDIA3-TID cell line compared to the control (**Fig S6D**). The ER specific localization of biotinylation in the TgPDIA3-TID cells was further confirmed through intracellular IFA with Avidin green staining (**Fig S6E**).

To increase specificity of binding to TgPDIA3, enrich for low abundance ER proteins, and control for highly expressed proteins from the cytosol, plasma membrane (PM), and other organelles, we conducted subcellular fractionation followed by a gradient centrifugation after biotin incubation (**Fig S6F**). The separation of the ER from the PM was validated by western blot analysis of gradient fractions, using PM and inner membrane complex (IMC) markers (αSAG1 and αGAP45, respectively) and ER markers (αTgSERCA and αTgPDIA3) (**Fig S6G**). ER (6a-7b) and cytosolic (S4) fractions were sent for LC-MS/MS analysis, together with equivalent fractions prepared by fractionation of the FBXO14-TID cell line [60], which expressed the cytosolic protein FBXO14 tagged with TurboID. The ideal control, a cell line expressing GFP alone in the ER, to account for non-specific interactions, could not be used as stable clonal lines expressing ER- localized GFP could not be successfully isolated, despite repeated attempts. To compensate for this limitation and increase specificity, we isolated fractions enriched in TgPDIA3 using an iodixanol gradient loaded with pre-enriched ER membranes (see schematic in Fig S6F). LC-MS/MS analysis revealed 56 proteins enriched in the TgPDIA3-TID ER fraction compared to the FBXO14-TID control. Both secretory and ER proteins were enriched in the TgPDIA3-TID ER vs. FBXO14-TID cytosolic, including TgSERCA and several microneme and rhoptry proteins (**Fig S6H-I**). Interestingly, there were 7 proteins that were significantly enriched in both the DVSF-IP and the TurboID-IP **(Table S3)**.

#### TgPDIA3 facilitates the secretion and maturation of micronemes and rhoptries

*T. gondii* microneme proteins begin their trafficking journey in the ER where they undergo initial processing before progressing through a series of maturation steps, ultimately localizing to the micronemes as fully mature MIC proteins [61]. Microneme secretion is triggered by a rise of cytosolic calcium, and ionophores, such as ionomycin or A32187, stimulate microneme secretion in the absence of host cells [11, 62]. Given that TgPDIA3 appears to interact with secretory proteins based on the biotinylation results, we next examined the microneme secretion and maturation profile of the *iΔTgPDIA3* (±ATc) mutant. We examined first both constitutive secretion of microneme protein 2 (MIC2) over 30 min and secretion induced by ionomycin for 3 min. In the *iΔTgPDIA3* (3 days +ATc) mutant, MIC2 secretion induced by ionomycin was significantly diminished compared to the *iΔTgPDIA3* (0 days with ATc) mutant or the *TatiΔku80* control. Secretion of dense granule protein 1 (GRA1), which is not dependent on calcium, served as secretion control (**Fig 4A and 4B**). This result indicates that TgPDIA3 plays a role in the pathway leading to calcium-induced microneme secretion.

**Fig 4.**
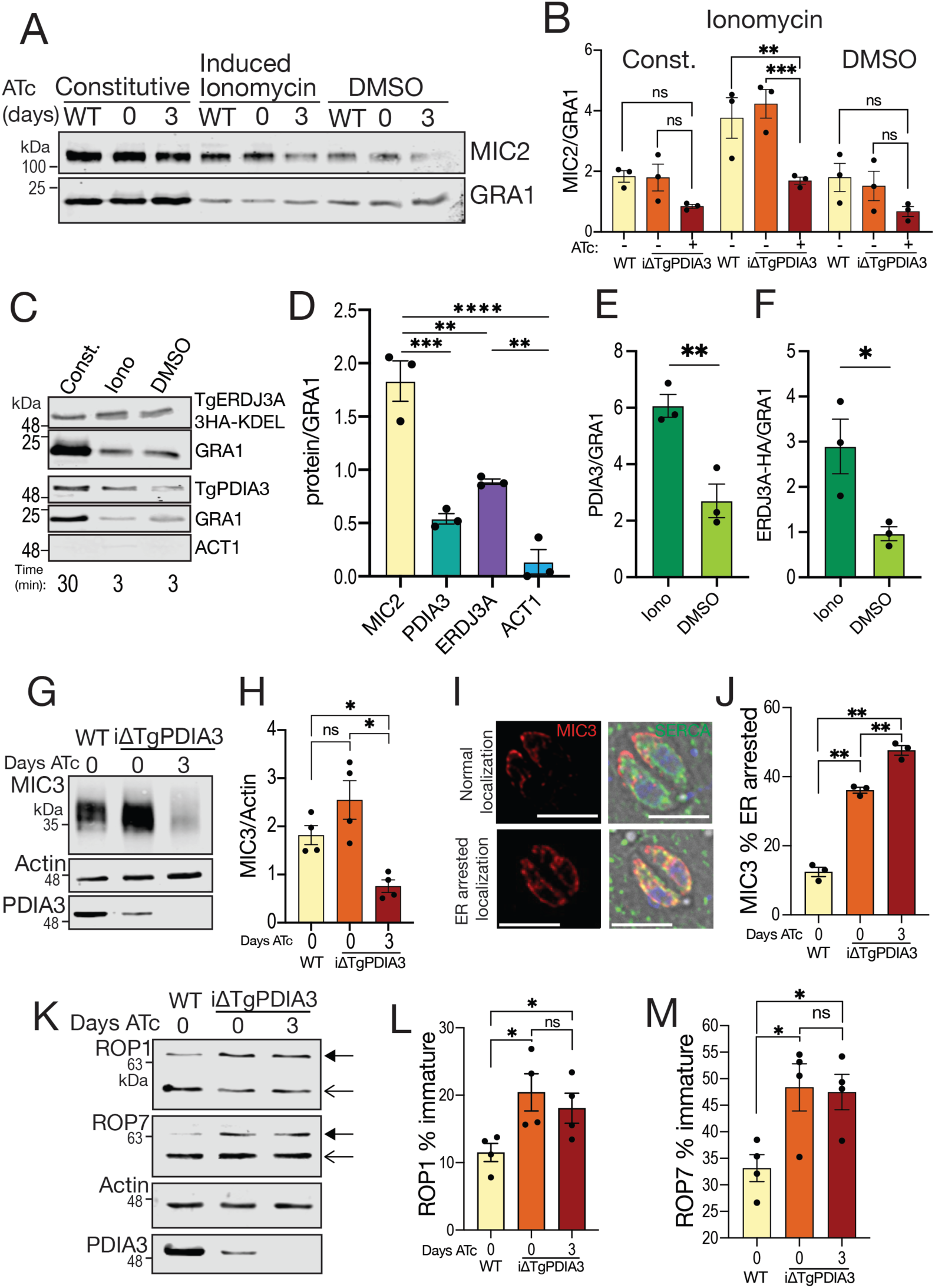
TgPDIA3 impacts microneme secretion and maturation, as well as rhoptry maturation. A) Representative western blots of microneme secretion assays of *iΔTgPDIA3* ± 2-day ATc pre-incubation including constitutive secretion (30 min) and induced secretion (3 min) with 1 μM ionomycin or DMSO (control). B) Quantification of standardized band intensity of αMIC2/αGRA1, two-way ANOVA for statistical analysis (n=3). C) Representative western blots of secretions of *TgERDJ3A*-*3HA-KDEL* or *TatiΔku80* including constitutive secretion (30 min) and induced secretion (3 min) with 1 μM ionomycin or DMSO (control) probed with αHA and αGRA1, or αTgPDIA3, αGRA1, and αACT1, respectively. D) Quantification of secreted proteins was performed by measuring the band intensity of each antibody signal relative to αGRA1, including αMIC2, αTgPDIA3, αHA, and αACT1. Statistical analysis was conducted using one-way ANOVA (n=3). E) Band intensity measurements of induced secretions were quantified as the ratio of αTgPDIA3 to αGRA1. Statistical significance was assessed using a Student’s *t*-test (*n* = 3). F) Band intensity measurements of induced secretions were quantified by measuring the ratio of αHA to αGRA1. Statistical significance was assessed using Student’s t-test (n=3). G) Representative western blot of *iΔTgPDIA3* ± 3-day ATc pre-incubation and *TatiΔku80* (WT) probed with αMIC3, αACT1, and αTgPDIA3. H) Quantification of band intensity ratio of αMIC3/αACT1 in *iΔTgPDIA3* ± 3-day ATc pre-incubation and *TatiΔku80* (WT). Statistical analysis was conducted using one-way ANOVA (n=4). I) Representative phenotype of MIC3 localization as observed by IFA and J) quantification of ER arrested localization of MIC3 (bar 5 μm). Statistical analysis was conducted using one-way ANOVA (n=3). K) Representative western blot probed with αROP1 and αROP7 in *iΔTgPDIA3* ± 3-day ATc pre-incubation and *TatiΔku80* (WT). Statistical analysis was conducted using one-way ANOVA (n=3). L) The percentage of immature protein was quantified by measuring band intensities of both forms (open arrows) detected with αROP1, with the immature form marked with a solid arrow. Data was analyzed by one-way ANOVA (n=4). M) The percentage of immature protein was quantified by measuring band intensities of both forms (open arrows) with the immature marked by a solid arrow, probed with αROP7. Data was analyzed by one-way ANOVA (n=4). All band intensities were measured using ImageStudioLite from LICORbio.

While performing microneme secretion assays we observed that TgPDIA3 was also secreted. To determine whether its secretion followed similar conditions as microneme proteins, we used the same protocol employed for measuring microneme secretion, and analyzed the presence of TgPDIA3 in the extracellular medium. Western blot analysis of the collected supernatant was performed using αTgPDIA3, with αGRA1 serving as a constitutively secreted control. Secretion of TgPDIA3 was analyzed by performing western blots of *TatiΔku80* supernatants with αTgPDIA3. We also tested another PDI, TgERDJ3A which was detected by probing a western blot of the *TgERDJ3A-3HA-KDEL* mutant supernatants with αHA. TgERDJ3A was constitutively secreted significantly more than actin, a control for parasite lysis (**Fig 4C-D**). Notably, the secretion of both TgPDIA3 and TgERDJ3A were stimulated by the calcium ionophore ionomycin (**Fig 4C, 4E-F**).

To further investigate the secretion of TgPDIA3 and TgERDJ3A, we conducted ultrastructure expansion microscopy (U-ExM) experiments to examine the presence of ER vesicles at the apical end and to determine whether TgPDIA3 localizes to the micronemes. We used αMIC2 and αTgPDIA3 to label the micronemes and the ER, respectively. Using Imaris (v9.0) microscopy image analysis software, we generated surface renderings based on fluorescence signals from αMIC2 and αTgPDIA3 staining. Our analysis did not reveal direct colocalization of micronemes (αMIC2) and the ER (αTgPDIA3); however, we observed numerous examples of proximity domains between the two (**supplemental video 1**). Proximity between the ER and micronemes would ensure that calcium is released precisely where it is needed for secretion.

*T. gondii* micronemes are rich in cysteine residues and disulfide bonds [63], the formation of which could be catalyzed by PDIs within the ER. To investigate the role of TgPDIA3 in the maturation of microneme proteins, specifically MIC3, which was enriched in both proteomic analyses, we performed western blots with whole cell lysates and probed them with αMIC3 to evaluate the changes in the ratios of mature and immature forms.

We found a significant reduction in total microneme protein 3 (MIC3) in the *iΔTgPDIA3* mutant cultured with ATc compared to controls (**Fig 4G-H**). To further assess the MIC3 processing defect, we conducted IFAs and observed two distinct patterns of its localization: an apical localization and an ER-arrested form with reduced apical localization. The *iΔTgPDIA3* (+ATc) mutant expressed higher level of the ER-arrested MIC3 compared to the control (*TatiΔku80*), indicating impaired trafficking of MIC3 (**Fig 4I-J**). Further support for the role of TgPDIA3 in MIC3 maturation, was provided by its enrichment in the +DVSF condition in the DVSF αTgPDIA3-IP (**Table 1**). TgPDIA3 disulfide bond formation in MIC3, likely stabilizes the protein in its correct conformation.

Considering the importance of rhoptry proteins for host cell invasion, PV formation, and host immune evasion [10], and their trafficking through the ER, we also explored TgPDIA3’s role in rhoptry maturation. Western blots analysis of rhoptry bulb proteins revealed significantly reduced levels of mature rhoptry bulb protein 1 (ROP1) and rhoptry bulb protein 7 (ROP7) in the *iΔTgPDIA3* mutant either with or without ATc compared to the parental control (*TatiΔku80*) (**Fig 4K-M**). Notably, rhoptry maturation was reduced in the *iΔTgPDIA3* (-ATc) mutant, which we attribute to partial down-regulation of *Tgpdia3* due to replacement of its native promoter with a weaker SAG4 promoter. This diminished expression, even in the absence of ATc, suggests that expression of TgPDIA3 is finely tuned and modest reductions can impact maturation of key secretory proteins (**Fig 4G and 4K**).

In summary, these findings suggest that TgPDIA3 is essential for the proper folding and maturation of microneme and rhoptry proteins in *T. gondii*. For microneme proteins, which are rich in disulfide bonds, TgPDIA3 likely facilitates disulfide bond formation necessary for their structural stability and function. For rhoptry proteins, TgPDIA3 may act indirectly through redox modulation of other proteins responsible for secretory protein processing, such as peptidases, which were enriched as redox interactors of TgPDIA3 in the DVSF αTgPDIA3 immunoprecipitation.

#### TgPDIA3 regulates TgSERCA and helps maintain low cytosolic calcium levels

The SERCA enzyme is a p-type ATPase that pumps calcium into the ER, helping to maintain low cytosolic calcium and high ER calcium levels, both of which are essential for proper cell signaling. Due to this critical role, SERCA activity is tightly regulated by both cytosolic and ER luminal proteins. In animals, PDIs interact with SERCA2b and IP_3_R through redox modifications. When ER calcium levels are high, the activity of SERCA2b (**Fig S7A**) is regulated by ERp57, which forms a disulfide bond with the cysteine residues present in the L4 loop of SERCA2b (**Fig S7B**), inhibiting its activity [31]. In animals, this interaction is enhanced by the binding of calreticulin or calnexin, to the ER luminal C-terminal tail of SERCA [40]. A modified version of the mammalian SERCA2b L4 luminal loop is found in TgSERCA, however, based on homology, the C-terminal calreticulin interacting tail [64] is absent. Notably, the L4 luminal loop of TgSERCA contains two cysteine residues at positions 947 and 967, mirroring the structure of the mammalian L4 loop in SERCA2b (**Fig S7**)

We first investigated the impact of TgPDIA3 in ER calcium regulation, by assessing cytosolic calcium responses to SERCA inhibitors in the *iΔTgPDIA3* (+ATc) mutant. Cytosolic calcium changes were measured using the genetically encoded calcium indicator (GECI) GCaMP6f (K_d_=375 nM), which allows assessment of calcium levels through fluorescence changes. We transfected the *iΔTgPDIA3* mutant with a gene expressing GCaMP6f-mScarlet and selected the clone with the highest dynamic range and lowest resting fluorescence [65]. Notably, upon addition of thapsigargin, a SERCA inhibitor, we observed a significantly greater increase in normalized fluorescence in cells cultured with ATc compared to the ones cultured without ATc, both in the presence or absence of extracellular calcium. These results support the notion of an increased calcium leakage from the ER when TgPDIA3 is absent (**Fig 5A-D**).

**Fig 5.**
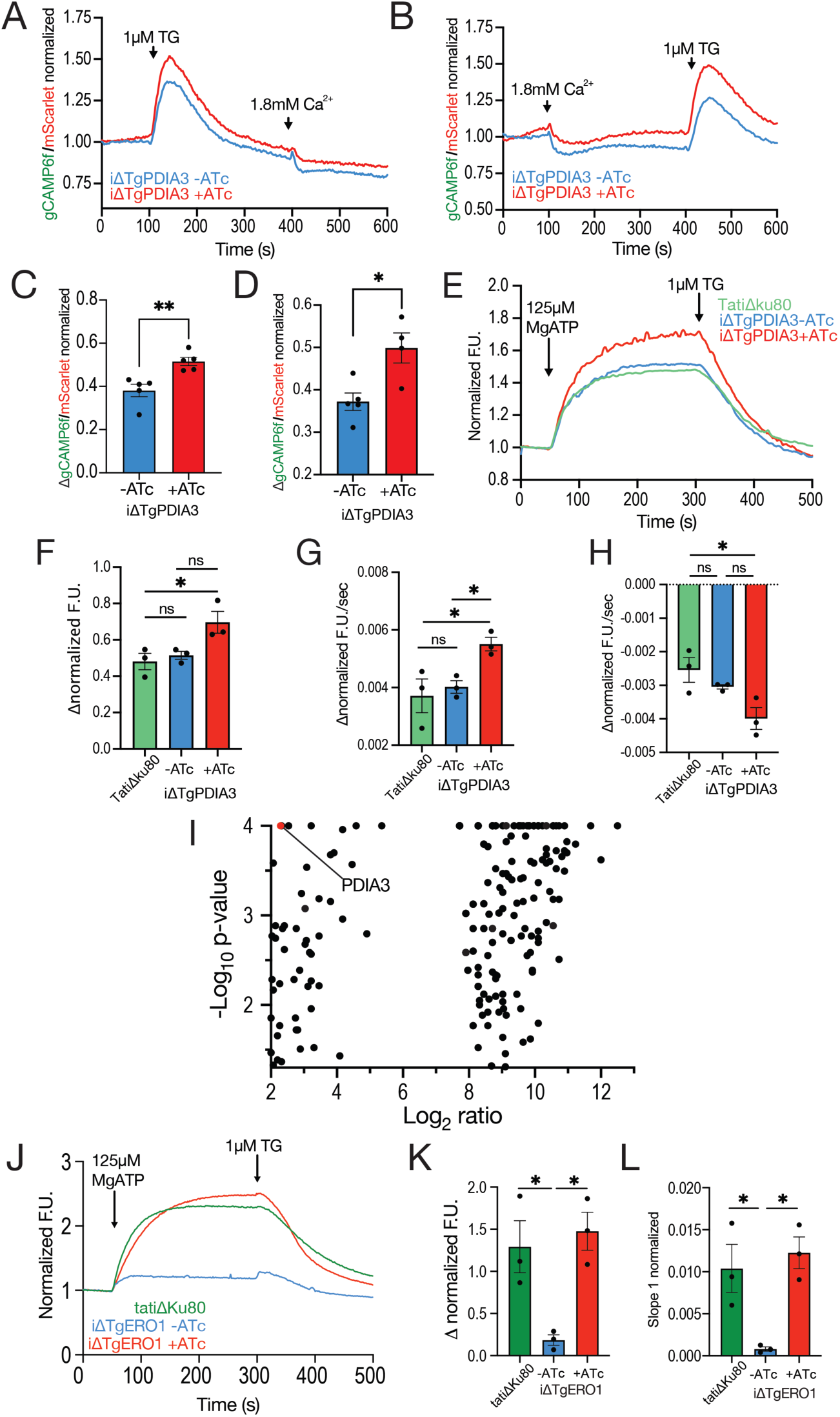
TgPDIA3 regulates TgSERCA. A) Average trace of normalized GCaMP6f/mScarlet fluorescence for cytosolic calcium measurements of the *iΔTgPDIA3* mutant ± 2-day ATc pre-incubation. 1 μM thapsigargin (TG) added at 100 s and 1.8 mM calcium added at 400 s. B) Average trace of normalized cytosolic GCaMP6f/mScarlet fluorescence for calcium measurements of the *iΔTgPDIA3* mutant ± 2-day ATc pre-incubation. 1.8 mM calcium was added at 100 s and 1 μM TG at 400s. C) Quantification of the change in fluorescence at 100 s. Data was analyzed using Student’s t-test (n=3). D) Quantification of the fluorescence change after 400s. Data was analyzed using Student’s t-test (n=3). E) Average traces of normalized Mag-Fluo-4 fluorescence of the *iΔTgPDIA3* mutant ± 2-day ATc pre-incubation compared to the *TatiΔku80* (control). F) Quantification of the total fluorescence increase after the addition of 125 μM MgATP. Statistical analysis was done using one-way ANOVA (n=3). G) Quantification of the slope at 50-100 seconds after the addition of 125 μM MgATP. Data was analyzed using one-way ANOVA (n=3). H) Quantification of the slope at 300-400 seconds after the addition of 1 μM TG. Data was analyzed using one-way ANOVA (n=3). I) Volcano plot showing DVSF-enriched hits for *TgSERCA-3HA* αHA-immunoprecipitation, with the Log_2_ ratio fold change (FC) on the x-axis and -Log_10_ of the p-value on the y-axis. Fisher’s exact test was used for statistical analysis (n=3). Proteins containing Trx-like domains are highlighted in red. J) Normalized fluorescence from MagFluo-4 traces from *iΔTgERO1-OE* mutant ± 2-day ATc pre-incubation compared to *TatiΔku80* (control). K) Quantification of the total increase in fluorescence after addition of 125 μM MgATP. Data was analyzed using one-way ANOVA (n=3). L) Quantification of the slope at 50-100 seconds after the addition of 125 μM MgATP. Data was analyzed using one-way ANOVA (n=3). The complete list of enriched peptides is in the supplemental Table: Proteomic Results.

To further explore this phenotype, we examined the potential role of TgPDIA3 in regulating SERCA activity. To this aim, we followed a well-established protocol for measuring ER-calcium uptake mediated by SERCA [66] that is validated by the specific block of uptake by thapsigargin. We loaded the *iΔTgPDIA3* mutant with the low affinity calcium indicator Mag-Fluo-4 (K_d_=22 μM) for an extended length of time to allow for its compartmentalization [48, 66]. Following digitonin permeabilization and washing, parasites were exposed to the SERCA substrate MgATP in the presence of 220 nM free calcium, and ER calcium uptake was measured. The resulting fluorescence increase specifically reflected calcium uptake by those organelles where SERCA is localized, primarily the ER and Golgi [67]. Further validation that this activity is due to SERCA calcium pumping comes from the effect of thapsigargin, which inhibits SERCA causing a decrease in MagFluo4 fluorescence due to calcium leakage from the ER. Upon MgATP addition to a suspension of the *iΔTgPDIA3* mutant (+ATc), we observed a significant increase in the change of normalized fluorescence compared to controls, *TatiΔku80* or *iΔTgPDIA3* (-ATc) (**Fig 5E-F**). This suggests that the knockdown of TgPDIA3 resulted in a significant increase in ER calcium uptake by TgSERCA, possibly due to the deficient regulation of TgSERCA at high ER calcium levels.

We also measured the rate of ER calcium uptake, which was significantly higher in the *iΔTgPDIA3* mutant (+ATc) (**Fig 5E and 5G**). After thapsigargin addition calcium leakage was both faster and greater in the absence of TgPDIA3 (**Fig 5E and 5H**). These findings suggest that TgPDIA3 may regulate SERCA activity which could be through the oxidation of its L4 luminal loop (**Fig S7**).

To further substantiate the possible regulation of TgSERCA by TgPDIA3, we performed αHA-immunoprecipitations with a 3xHA-tagged TgSERCA, *iΔTgSERCA-3HA*, in the presence of DVSF. LC-MS/MS analysis of the samples revealed significant enrichment of TgPDIA3 in the +DVSF condition (**Fig 5I**). Collectively, these data suggest that TgPDIA3 plays a regulatory role in TgSERCA function likely through redox modifications of TgSERCA’s L4 luminal loop.

Considering this finding and the fact that SERCA2b is regulated by multiple redox proteins in animals, we sought to explore whether other ER redox-related proteins might also regulate TgSERCA. We considered TgERO1, which was highly enriched in the LC-MS/MS analysis of the αTgPDIA3 +DVSF sample (**Fig 3D**). To investigate the role of TgERO1 in SERCA activity, we used the *iΔTgERO1-3Ty* mutant. Interestingly, we observed an 80% decrease in SERCA activity in the *iΔTgERO1-3Ty* mutant (-ATc). The activity was recovered to the levels of the *TatiΔku80* control when the *iΔTgERO1-3Ty* mutant was pre-incubated with ATc (**Fig 5J-L**). Previous studies have shown that ERO1 overexpression, especially in the absence of SEPN1, can lead to excessive production of H_2_O_2_ in the ER. This causes hyperoxidation of the L4 luminal loop of TgSERCA, severely decreasing its activity [68]. RNA-seq datasets from toxodb.org show that *Tgero1* (TGME49_300380) is expressed at 52 TPM (transcripts per million) in tachyzoites, whereas *sag4* (TGME49_280570), from which the promoter domain was used for the generation of the *iΔTgERO1-3Ty* line, is expressed at 1692 TPM [69]. This represents a 32-fold higher transcription level compared to endogenous *Tgero1*, suggesting that Tgero1 is overexpressed in the *iΔTgERO1-3Ty* mutant, explaining the observed loss of SERCA activity. TgERO1 was not enriched in the +DVSF condition of the *iΔTgSERCA-3HA* αHA-immunoprecipitation supporting that the reduction of SERCA activity was not through direct disulfide exchange with ERO1.

The ER is one of the main sites of ATP consumption, and in mammals, some redox proteins are active at Mitochondria-Associated Membranes (MAMs) to regulate calcium transfer from the ER to the mitochondria in order to increase ATP production or induce apoptosis during prolonged ER stress [29]. Since there is a defect in ER calcium regulation in the *iΔTgPDIA3* mutant we next explored the potential role of TgPDIA3 in calcium transfer from the ER to the mitochondrion. To test this, we transfected the *iΔTgPDIA3* mutant with a GCaMP6f-SOD2 plasmid which allows the expression of GCaMP6 in the mitochondrion [65, 70, 71]. Since the *T. gondii* mitochondrion does not have direct access to extracellular calcium [48], and instead relies on calcium accumulation in ER-mitochondrion microdomains, we used thapsigargin, to stimulate calcium leakage from the ER. The resulting calcium transfer into the mitochondrion was detected as an increase in the GCaMP6 fluorescence. This increase was lower in the *iΔTgPDIA3* mutant, although the difference was not statistically significant. These findings suggest a potential defect in the ER to mitochondria transfer of calcium in the mutant (**Fig S8A-B**). We also used zaprinast, a phosphodiesterase inhibitor which ultimately causes the release of calcium from the ER and acidic organelles. In the *iΔTgPDIA3* mutant (+ATc) there was significantly less calcium taken into the mitochondria compared to the control (-ATc) (**Fig S8C-D**). A similar trend was observed following the addition of glycyl-L-phenylalanine 2-naphthylamide (GPN), a lysomotrophic agent that mobilizes calcium from acidic stores [72] like the plant-like vacuolar compartment or PLVAC, [73, 74], an acidic compartment with lysosomal and secretory functions (**Fig S8E-F**). This result was unexpected given that SERCA is not properly regulated in the *iΔTgPDIA3* mutant, and more calcium is being pumped into the ER, we would have anticipated either no change or an increase in calcium transfer to the mitochondrion. One possible explanation is that the ER leaks calcium in a more diffuse manner rather than specifically at ER-mitochondria contact sites, resulting in greater calcium dilution in the cytosol. This hypothesis is supported by the cytosolic calcium response shown in . In summary, our results indicate that TgPDIA3 regulates TgSERCA activity, which may, in turn, influence calcium levels in the cytosol, ER, and mitochondrion.

## Discussion

In this study, we characterized several redox proteins of the *T. gondii* ER including TgPDIA3 (TGGT1_211680), TgPDIA6 (TGGT1_249270), TgERO1 (TGGT1_300380), and TgERDJ3A (TGGT1_209950). We demonstrated the protein folding ability of TgPDIA3 and TgPDIA6 and established the role of TgPDIA3 in microneme secretion, as well as the maturation of MIC3 and rhoptry proteins. Additionally, we characterized TgPDIA3 as a catalytically active protein disulfide isomerase and identified several of its substrates. Both TgPDIA3 and TgERO1 influenced the calcium-pumping activity of TgSERCA, which was further supported by the demonstration of TgPDIA3-SERCA interaction. Furthermore, the redox activity of TgPDIA3 impacted ER calcium homeostasis, subsequently affecting calcium transfer to the mitochondrion (**Fig. 6)**.

**Fig 6.**
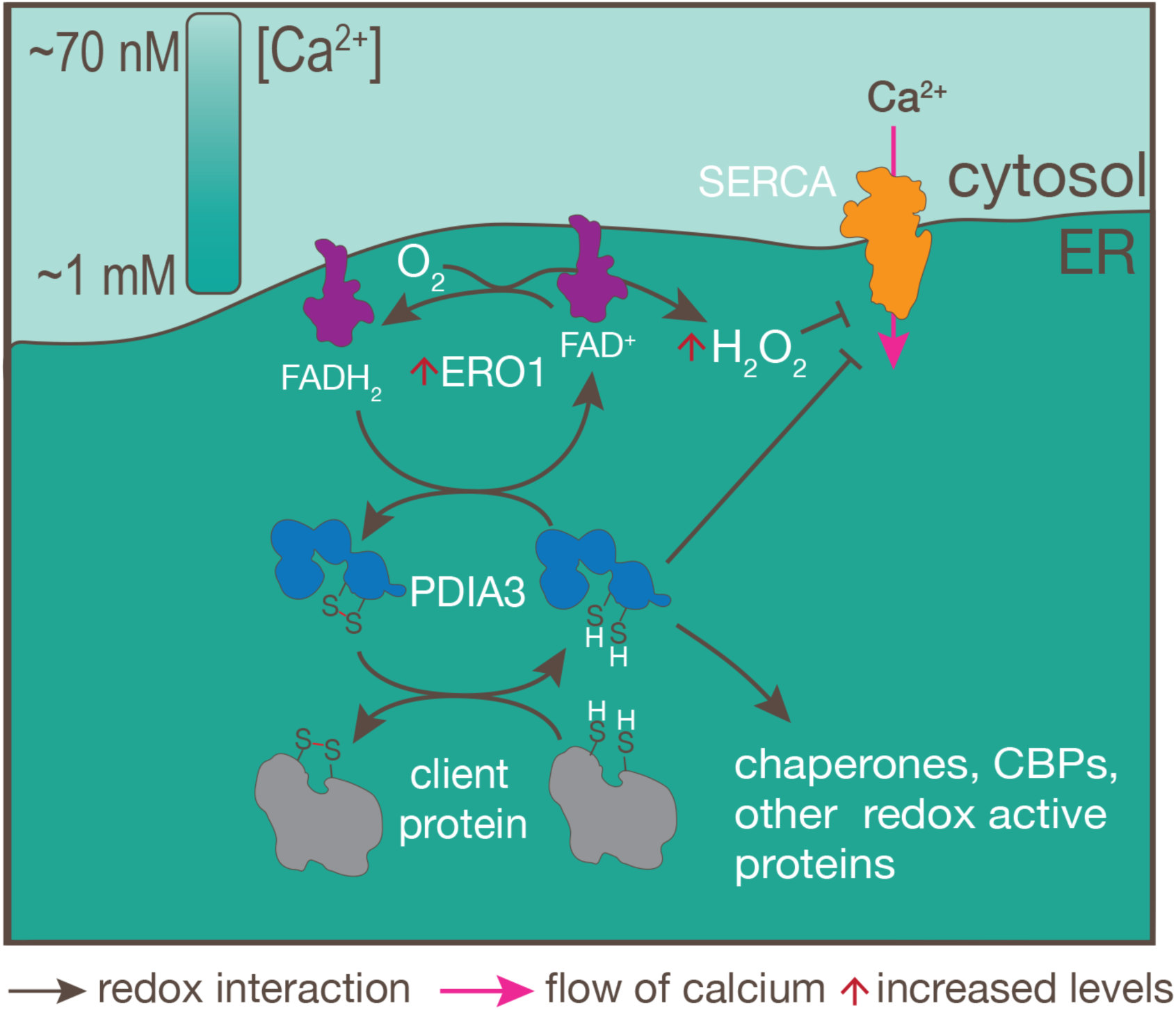
Model of the role of TgPDIA3 in the ER. TgPDIA3 interaction with client proteins and its catalytic activity of disulfide bond formation. Oxidation of TgPDIA3 by TgERO1 re-activates it, and as a result TgERO1 produces hydrogen peroxide. Redox regulation of TgSERCA by TgPDIA3 during high ER calcium and by ROS upon ERO1 over expression is shown. Other interactions of TgPDIA3 are also possible based on the results of the DVSF IP enriched proteins.

Denatured GFP serves as an effective substrate for testing the refolding activity of PDIs, with fluorescence readings providing a practical measure of refolding efficiency [45]. Using this approach, we demonstrated that both recombinant TgPDIA3 and TgPDIA6 exhibit refolding activity, facilitating the recovery of denatured GFP. Although GFP contains two cysteine residues, it does not naturally form disulfide bonds [75], making it unlikely that the observed effect is mediated by disulfide bond formation. Instead, this result could reflect the chaperoning abilities of TgPDIA3 and TgPDIA6. PDIs may act as chaperones through interactions with hydrophobic residues in its b and b’ domains, which have been shown *in vitro* to reduce aggregation of unfolded proteins [76], promoting protein folding. Further experiments are needed to confirm this possibility.

Previous studies have used DVSF to capture redox interactions between PDIs and their substrates in Apicomplexans [77]. Using a DVSF αTgPDIA3-IP approach, we identified numerous potential redox partners of TgPDIA3, including other PDIs, suggesting that *T. gondii* PDIs may interact with each other similarly to their mammalian counterparts [58, 59]. The enrichment of TgERO1 in the DVSF αTgPDIA3-IP provides a potential mechanism for PDI cysteine re-oxidation, similar to mammalian PDIs, where ERO1 likely oxidizes disulfide bonds on PDI to sustain its catalytic function [57, 78].

Interestingly, metallopeptidases were among the most significantly enriched proteins, which represents a previously unreported interaction that may indicate a regulatory role for TgPDIA3 in peptidase activity. However, further studies are needed to determine the functional significance of this interaction. Metallopeptidases have been previously studied in *T. gondii* [79–82] and most characterized members to date have been shown to be secreted, with a hypothesized role in cleaving host proteins. We also observed enrichment of Cdc48 suggesting a possible involvement of TgPDIA3 in endoplasmic reticulum-associated degradation (ERAD) [83]. Additionally, several proteins predicted as glycolsyltransferases were identified, aligning with the known function of mammalian PDIA3 in catalyzing disulfide bond formation in glycoproteins. Future work on these TgPDIA3 substrates is warranted to better understand their functional significance in *T. gondii* biology. Understanding the molecular mechanisms of these interactions could reveal new insights into redox regulation, protein folding, and ER-associated processes in the parasite.

Apicomplexans are named for their apical complex, a critical assembly of structures and organelles that secrete effectors essential for the various phases of the parasite’s lytic cycle [7, 8]. Micronemes and rhoptries, specialized secretory organelles located within the apical complex, play crucial roles in the invasion of host cells [84]. *T. gondii* secreted proteins from micronemes and rhoptries go through multiple modifications in various subcellular compartments before reaching their mature forms [85]. TgPDIA3-TID LC-MS/MS results showed an enrichment of secreted proteins, suggesting that TgPDIA3 interacts closely with *T. gondii* secreted effectors, likely as they are trafficked through the ER [61]. In this regard, a recent study about the characterization of TgPDIA3 (aka TgPDI8) also found that it was proximal to secretory proteins [86]. Our study found that TgPDIA3 is important for the maturation of select microneme and rhoptry proteins, including MIC3, ROP1, and ROP7. We hypothesize that TgPDIA3 facilitates this maturation through disulfide bond formation, especially for MIC3, which was significantly enriched in the DVSF αTgPDIA3-IP. For rhoptry proteins, which contain fewer disulfide bonds and which were not enriched as redox substrates of TgPDIA3, the role of TgPDIA3 in their maturation could be more indirect, potentially through the redox modulation of other proteins such as peptidases. Interestingly, several peptidases were enriched as redox interactors of TgPDIA3 in the DVSF αTgPDIA3-IP. Since parasite invasion, gliding motility, and egress all depend on microneme secretion, and this secretion is tightly regulated by intracellular calcium signaling [11, 62], TgPDIA3 may also contribute to these processes indirectly. As the largest intracellular calcium store, the ER is responsible for calcium distribution, and forms connections with other calcium stores like the mitochondrion and the PLVAC [48]. The ER of *T. gondii* forms a vast, interconnected tubular network that extends throughout the entire parasite [87]. Notably, U-ExM experiments revealed close proximity between the ER and micronemes (**Supplemental Video 1**), suggesting an alternative possible role in calcium release from the ER in domains close to the micronemes to facilitate calcium induced microneme secretion. The presence of calcium microdomains between the ER and other organelles is well established [88, 89], and serves in preventing global cytosolic calcium increases that could lead to cytotoxicity or unintended signaling. Since microneme proteins undergo maturation in a series of vesicular compartments exiting the ER and trafficking through the Golgi [61], their primary transport and maturation would not depend on ER-microneme proximity. However, ER-microneme proximity could play a role in the localized release of calcium from the ER, which may be important for microneme secretion. This spatial arrangement may provide an advantage to the parasite by enabling rapid and localized calcium release to micronemes without the need for global cytosolic calcium elevation. In this regard, our study showed that downregulation of TgPDIA3 resulted in a reduction of calcium-induced microneme secretion while constitutive secretion showed no significant difference. The observed reduction in SERCA regulation in the *iΔTgPDIA3* (+ATc) mutant further supports our hypothesis as the increased activity of TgSERCA would result in a reduction in the ER-microneme microdomain calcium concentration. These results would also explain the invasion defect observed in the *iΔTgPDIA3* mutant.

In mammals, some ER-resident proteins, such as BiP and calreticulin, have been shown to be secreted during calcium-induced ER stress [90]. Calreticulin has been proposed as an ’eat-me’ signal in dead or dying cells and can also promote the phagocytic uptake of cells undergoing ER calcium depletion [91]. Additionally, impaired PDI secretion from injured endothelial cells has been linked to defective blood clot formation [92, 93]. HsPDIA3, specifically, has been associated with extracellular matrix mineralization through its interaction with bioactive vitamin D and bone morphogenetic protein-2 (BMP2) [94, 95]. Similarly, we found that both TgPDIA3 and TgERDJ3A were secreted in a calcium dependent manner, like micronemes. Previous studies have suggested the potential use of TgPDIA3 as a diagnostic tool for *T. gondii* infection based on its ability to be recognized by IgA antibodies in human tears and milk [96]. Based on its abundance, potential for secretion, and preliminary findings TgPDIA3 may also represent a promising candidate for *T. gondii* vaccine development [97].

Previous studies have demonstrated that calcium movements in the mammalian endoplasmic reticulum is redox-regulated through cysteine modifications of various ER membrane proteins involved in calcium transport [30, 31]. Our lab previously showed that TgSERCA is essential for replenishing ER calcium during the lytic cycle and facilitates its transfer to other compartments, ensuring the replenishment of all intracellular calcium stores [48].

In the *iΔTgPDIA3* mutant, we observed increased ER calcium uptake most likely through the activity of TgSERCA, followed by accelerated calcium leakage after SERCA inhibition with thapsigargin, suggesting a regulatory role of TgPDIA3 in TgSERCA function [31]. We hypothesize that this regulation occurs through direct redox modification of TgSERCA’s L4 luminal loop by TgPDIA3 under high ER calcium conditions. Supporting this hypothesis, TgPDIA3 was enriched in the DVSF *TgSERCA-HA* αHA-IP. Additionally, TgERO1 overexpression led to a dramatic reduction in SERCA activity, likely due to hyperoxidation of the TgSERCA’s L4 loop caused by excessive H₂O₂ production from TgERO1 [68]. We noticed that overexpression of TgERO1 in the *i11TgERO1* (-ATc) mutant was not fitness conferring for the parasite even though TgSERCA is essential for parasite survival [48]. We propose two hypotheses to explain this result. First, the residual SERCA activity in the *i11TgERO1* (-ATc) mutant may be sufficient to support normal intracellular growth. Alternatively, while SERCA may function normally during intracellular replication, the stress of parasite purification and exposure to extracellular conditions could trigger elevated H₂O₂ production by overexpressed ERO1, leading to a sharp inhibition of SERCA activity. Future experiments will focus on measuring of H_2_O_2_ in extracellular *i11TgERO1* (-ATc) parasites, comparing conditions with and without ATc to assess the impact of ERO1 overexpression.

It is interesting that despite increased calcium leakage into the cytosol, mitochondrial calcium uptake was diminished rather than elevated. PDIs are known to localize to mitochondrial-associated membranes (MAMs) [98] and may facilitate calcium transfer from the ER to the mitochondrion. Our experiments showed reduced mitochondrial calcium uptake in the *iΔTgPDIA3* (+ATc) mutant, which we hypothesize is due to a diffuse calcium leakage from the ER after inhibition of SERCA rather than being localized to specific microdomains between the ER and mitochondrion. It is possible that TgPDIA3 may be important for the localization of specific proteins involved in the transfer of calcium between the ER and the mitochondria. More work is needed to characterize this phenomenon.

In conclusion, our study highlights the critical role of TgPDIA3 in the redox regulation of calcium signaling in *T. gondii*. By modulating ER calcium homeostasis, TgPDIA3 influences key processes such as microneme secretion, which is essential for parasite invasion. We also identified several TgPDIA3 substrates, and demonstrated its involvement in parasite replication, and the maturation of secreted proteins. These findings represent the first evidence of the link between redox regulation and calcium homeostasis in *T. gondii*, expanding our understanding of the role of the *Toxoplasma* ER in calcium storage and signaling. Calcium signaling is crucial for the progression of the lytic cycle [99–101], with intracellular stores playing a key role in this process [48, 102]. Further studies are needed to elucidate the mechanisms by which the ER regulates calcium storage and release.

## Materials and Methods

### Phylogenetic Analysis

Sequences were obtained through the NCBI database and VEUpath DB [103]. Sequences were aligned in MEGA-X using Clustal W with manual trimming. Maximum likelihood trees were also constructed on MEGA-X [104]. Phylogeny test was conducted using the bootstrap method with 1000 bootstrap replications. Amino acid substitution using the Jones-Taylor-Thornton (JTT) Model. Uniform Rates, and all sites were used in tree construction. The Nearest-Neighbor-Interchange (NNI) maximum likelihood heuristic model was used.

#### Cell Culture

Human Telomerase reverse transcriptase (hTERT) fibroblasts were cultured in Dulbecco’s modified Eagle medium (DMEM-HG) with 10% bovine calf serum (BCS) at 37°C with 5% CO_2_. Human Foreskin Fibroblasts (HFF) cells were cultured in DMEM-HG with 15% FBS at 37°C with 5% CO_2_. *T. gondii* tachyzoites were cultured in hTERT fibroblasts in DMEM-HG with 1% BCS at 37°C with 5% CO_2_. C-terminally tagged mutants were grown in media supplemented with 6.8 µg/mL chloramphenicol and promoter insertion mutants were grown with 1 µM pyrimethamine. When needed for down-regulation of target genes, parasites were grown with 0.5 µg/mL anhydrous tetracycline.

#### Generation of Plasmids and Mutants

Guide RNAs (gRNAs) were added into Cas9s on primers using Q5 PCR and site-directed mutagenesis as previously described [105]. Plasmids with modified retention signals were also altered using Q5 PCR and site-directed mutagenesis. Recombinant protein plasmids were created using Gibson assembly [106]. All inserted sequences were confirmed with sanger sequencing.

Genes were tagged using a CRISPR/Cas9 approach. To create all mutants, *TatiΔku80* parasites were transfected with a Cas9 plasmid containing a gRNA specific to either the 5’UTR or 3’UTR just outside the gene of interest (GOI). Corresponding repair templates were amplified with homology just inside the GOI and on the opposite side of the gRNA to introduce a regulatable promoter and/or tag at the 5’ end of a GOI or a C-terminal tag at the 3’ end of a GOI. Promoter insertion and tagged mutants were enriched with drug selection, subcloned and validated genomically via PCR and protein tagging was validated via IFA and western blot. Fluorescent mutants, that were created by transfection of tdTomato, GCaMP6f-mScarlet, or linearized GCaMP6f-SOD2 [65], were enriched through fluorescence sorting on a BioRad S3 cell sorter, and the fluorescence of parasites in the mixed populations were validated via live cell imaging on a DeltaVision, DVElite. All mutants were subcloned to obtain single population clones after multiple rounds of enrichment. To select GCaMP6f clones, the fluorescence of 2x10^7^ parasites was measured on a BioTek Synergy H1 Hybrid Reader and 1 µM ionomycin was added to determine calcium dynamic range. Clones with the lowest resting fluorescence and the largest dynamic range were selected and used for the experiments.

#### Gene cloning and protein purification

*Tgpdia3* (TGGT1_211680) and *tgpdia6* (TGGT1_249270) were amplified from *T. gondii* RH cDNA, and GFP was amplified from pTREX-GFP plasmid, and cloned into the pQE-80L vector by Gibson Assembly, a 5’ exonuclease-3’ extension method of DNA assembly, and transformed into NEB 5α competent *E. coli.* For each gene, 2 of these bacterial colonies were sent for sequencing for confirmation of proper assembly. The plasmids were then transformed into protein-expressing bacteria, either BL21 DE3 or BL21 codon+ *E. coli,* and colonies were tested for IPTG induced gene expression. Gene expression and protein solubility was optimized by altering IPTG concentrations and induction temperatures and times. Protein induction and solubility was visualized on a coomassie stained gel (**Fig S3B and S5A-B**) and western blots of the induced and uninduced samples were probed with α-His antibody at 1:1,000 to better determine solubility of the proteins. Soluble proteins were purified using a HisPur Ni-NTA Chromatography Cartridge [CAT:90098].

#### Generation of antibodies in mice

The antigen mixture was prepared using the purified recombinant protein described above and either Freund’s Complete adjuvant [CAT: F5881] (for initial injection) or Freund’s Incomplete adjuvant [CAT: F5506] (for subsequent booster injections). Mice were interperitoneally injected with the antigen once every 2 weeks for a total of 4 injections, 100 µg of protein per mouse was used for the primary inoculation and 50 µg of protein per mouse was used for the boosts. Serums were collected after each injection to test for the presence of antibodies before sacrificing mice. Mice were sacrificed and bleeds were stored at 4°C over night to allow for separation of serum and red blood cells [51, 107].

#### IACUC statement

Work with mice was carried out in strict accordance with the Public Health Service Policy on Humane Care and Use of Laboratory Animals and Association for the Assessment and Accreditation of Laboratory Animal Care guidelines. The animal protocol was approved by the University of Georgia’s Committee on the Use and Care of Animals (protocol A2021 03-005-A5). All efforts were made to humanely euthanize the mice before final blood collection.

#### Immunofluorescence Assays

Immunofluorescence assays were conducted as previously described [107]. Tachyzoites were grown on confluent HFF or hTERT cells on coverslips for ∼24 hours. Cells were either fixed and permeabilized with 100% methanol for 2 minutes followed by rehydration with PBS or fixed with 3% paraformaldehyde and permeabilized with 0.25% Triton X-100 in PBS. Coverslips were blocked and antibody stained in 3% bovine serum albumin (BSA) in PBS pH 8.0. Coverslips were washed with PBS pH 8.0 after blocking, primary staining (mouse-Ty 1:200 (from Drew Ethridge) and rat αHA at 1:50 (Roche)) and secondary staining. Cells were imaged using a DeltaVision, DVElite.

#### Western Blots

For general protein lysis, parasites were incubated with Cellytic M [CAT:C2978] (with DNAse and RNAse) for 5 minutes at RT. Proteins were then boiled with Laemmli BME for 5 minutes. For western blots, SDS-PAGE separated proteins were transferred to a nitrocellulose membrane. Membranes were blocked with 5% milk in PBS-T before incubating with primary antibodies in PBS-T followed by secondary antibodies in PBS-T, blots were washed with PBS-T in between steps and before imaging. Blots were imaged using a LiCOR Odyssey CLx imager.

#### Plaque Assay

Plaque assays were performed as previously described [108]. Freshly egressed tachyzoites were collected, filtered, and counted. 200 tachyzoites/well were used to infect confluent hTERT monolayers in 6-well plates and incubated at 37°C for 7 days. Mutant and control strains were incubated with or without ATc, after which monolayers were fixed for 5 minutes with 100% ethanol, stained with crystal violet for 5 minutes, and washed with PBS. Plates were imaged and plaque sizes were measured using FIJI [109]. 16 plaques were measured per biological replicate and 3 biological replicates were used per condition.

#### Growth and Replication Assays

Growth assays were performed as previously described [110]. Confluent monolayers of hTERT cells in a 96-well plate were infected with freshly egressed tdTomato expressing tachyzoites at 4000 parasites per well. Mutants were incubated either with or without ATc and fluorescence was measured daily on a BioTek Synergy H1 Hybrid Reader for 8 days to track parasite growth. A standard curve was created using serial dilutions of fluorescent parasites in order to calculate the number of parasites per well based on their fluorescence level.

Replication assays were conducted as previously described [107] with a few modifications. 5x10^5^ freshly egressed RFP expressing tachyzoites were used to infect confluent hTERT fibroblast monolayers on coverslips. Coverslips were fixed and mounted 20 hpi. The number of 1, 2, 4, and 8+ parasite containing vacuoles were counted until >100 total vacuoles were counted per condition and quantified.

#### Invasion (Green/Red) Assay

Invasion assays were conducted as previously described [50] with a few modifications. 2x10^7^ of freshly egressed RFP expressing tachyzoites were washed and resuspended in invasion media (3% FBS, 10 mM HEPES, in DMEM-HG, pH 7.4) and allowed to settle on HFF coated coverslips for 20 minutes on ice. Plates of parasites were placed in a 37°C bath for 5 min to stimulate invasion and then returned to ice and immediately fixed with 3% paraformaldehyde to stop invasion. Coverslips were blocked and stained with rabbit αSAG1 at 1:1000 (a gift from Vern Carruthers) to differentiate the extracellular parasites from the intracellular ones. Parasites were then counted, and the number of red parasites (all) and green (extracellular) was obtained and graphed as a percentage of total parasites that invaded.

#### In vitro protein folding assays

To evaluate the protein chaperoning abilities of TgPDIA3 and TgPDIA6, we adapted a protein chaperone assay utilizing recombinant TgPDIs and a recombinant acid-denatured GFP [53]. To begin all proteins were diluted and concentrated in PBS to remove contaminating imidazole from the purification process (because it has chaperoning abilities on its own). Recombinant GFP was diluted in denaturing buffer (0.3 mM EDTA, 50 mM Tris-HCl, pH 7.5) and mixed with an equal volume of 125 mM HCl to acid-denature GFP for 1 minute at RT. Denatured GFP was added to a cuvette of renaturation buffer (25 mM MgCl_2_, 100 mM KCl, 50 mM Tris-HCl, pH 7.5) either with or without PDI after 20 sec of reading. Re-folding was measured as a gain in fluorescence using a Hitachi F-7000 Fluorescence Spectrophotometer [53]. Spontaneous refolding in the absence of PDI was compared to refolding in the presence of different concentrations ratios of GFP to PDI.

#### DVSF Crosslinking

DVSF crosslinking was conducted as previously described [77] with a few modifications. In brief, the chemical crosslinker Divinyl Sulfone (DVSF), 97% [CAT: L12827-09] was used to covalently bind PDI’s catalytically active cysteines in the CXXC motif with cysteine residues in client proteins during disulfide bond exchange. DVSF was diluted at a concentration of 3 µL DVSF/10 mL BAG and freshly collected parasites were resuspended in 1 mL of the BAG with DVSF or BAG without DVSF and incubated in a 37°C water bath with rotation for 30 minutes. Parasites were then centrifuged and washed once with BAG before centrifugation and resuspension in lysis buffer. Parasites were lysed and proteins collected for western blot analysis. For enrichment of membrane proteins, cells were subjected to 3 freeze-thaw cycles and lysates were centrifuged to separate soluble (supernatant) and membrane (pellet) proteins. Proteins were then solubilized using 1% SDS or 1% NP-40 for subsequent analysis.

#### TgPDIA3 co-Immunoprecipitation

The IP was carried out as previously described with some modifications [111]. Protein G magnetic beads [CAT: 88847] were gently vortexed for resuspension and 100 µL of beads was washed, collected and incubated with mouse αTgPDIA3 in 1 x coupling buffer using 20 x coupling buffer (200 mM NaH_2_PO_4_, 3 M NaCl, pH 7.2) and 5% fresh Lysis buffer (150 mM NaCl, 20 mM Tris pH 7.6, 0.1% SDS, 1% Triton X-100). The antibodies were crosslinked to the beads using 220 mM DMP [CAT: D8388] in 200 mM sodium borate pH 9.0. The coupling was quenched using 200 mM ethanolamine, pH 8.5. Parasites were collected as previously described for DVSF crosslinking and lysed for 5 minutes on ice with Lysis buffer with protease inhibitor [CAT: 11836170001] with periodic vortexing to facilitate lysis. Lysates were centrifuged at 21,000 x g for 5 minutes at 4°C to clarify the lysates and the supernatant was added to the collected beads and incubated at 4°C overnight. The unbound protein was then collected, and the beads were washed with the lysis buffer 3 times before boiling with 1 x Laemmli for western blot testing or the beads were frozen dry and sent for LC-MS/MS analysis.

### TgSERCA-3HA co-Immunoprecipitation

Parasites were collected as previously described for DVSF crosslinking including the enrichment of membrane proteins step. After freeze-thaw, initial resuspension of pellets, and removal of soluble proteins, the pellets enriched with membrane proteins were lysed for 5 minutes on ice in lysis buffer (50 mM HEPES pH 7.4, 50 mM NaCl, 1% NP-40, and mini complete protease inhibitor [CAT: 11836170001]) with periodic vortexing to facilitate lysis. Lysates were centrifuged at 21,000 x g for 5 minutes at 4°C. After washing 30 µL of Pierce α-HA magnetic beads (Cat. No. 88836) three times with lysis buffer, clarified lysates from 4 × 10⁸ parasites were added to the beads and incubated overnight at 4 °C. The beads were collected, and unbound proteins were removed. The beads were then washed with the lysis buffer 3 times before boiling with 1 x Laemmli in lysis buffer for western blot testing or the beads were frozen dry and sent for LC-MS/MS analysis.

#### Subcellular fractionation and Gradient centrifugation

Subcellular fractionation was carried out as previously described [73, 112]. Following a 1 hr incubation with 50 μM biotin at 37**°**C, parasites were collected, needle lysed out of host cells, and filtered through an 8 µm nuclepore membrane [73]. Parasites were then washed with BAG, counted, and lysed by grinding with silicone carbide on ice for 30-sec intervals with 30-sec pauses in between for a total grinding time of approximately 2 min, in order to keep organelles intact. Parasites and silicon carbide were resuspended in lysis buffer (50 mM KCl, 4 mM MgCl_2_, 0.5 mM EDTA, 20 mM HEPES-KOH pH 7.2, 125 mM sucrose, mini complete protease inhibitor, 12 µg/mL DNase, 12 µg/mL RNase, and 8 µg/mL nocodazole). Silicon carbide was removed through a series of low-g centrifugations. Fractions were collected after each centrifugation step (**Fig S6F**). Pellets and supernatants were collected, and supernatants re-spun for the following centrifugation until pellet (P3) fraction, which was homogenized and mixed with the 20% layer of an Optiprep [CAT: M1248-250] gradient for a total volume of 12 mL (**Fig S6F**). The gradient was top loaded and then centrifuged at 50,000 x g for 1 hour using a SW-41Ti rotor in an Optima XE-100 Ultracentrifuge. 24-500 μL fractions were collected from the top of the gradient and labeled as (starting from the top) 1a, 1b, 2a, 2b, etc. until 12b (the bottom fraction). Fractions of interest were diluted fivefold, re-centrifuged at 100,000xg for 1 hr, and resuspended in lysis buffer and protease inhibitor to remove potentially contaminating iodixanol from the Optiprep.

#### Streptavidin co-immunoprecipitation

Fractions of interest from both *TgPDIA3-TID-3HA-GEEL* and *FBXO14-TID* were lysed in RIPA buffer (150 mM NaCl, 0.1% SDS, 0.5% sodium deoxycholate, and 1% NP-40 in 50 mM HEPES pH 7.5) as previously described [113] and incubated with Streptavidin magnetic beads for 1 hr at RT. Proteins were eluted from the beads using a buffer with excess biotin (20 mM Biotin 1% SDS and 25 mM Tris, pH 7.4) at 75**°**C for 30 min. Co-IP eluates were sent to the University of Nebraska Proteomics & Metabolomics Facility for LC-MS/MS analysis. 2 biological replicates each for S4 and the combined ER gradient fractions 6a, 6b, 7a, and 7b were sent. Results were analyzed using Scaffold to determine those proteins enriched in the ER fractions in TgPDIA3-TurboID cell line compared to S4 in FBXO14 cytosolic controls.

#### Microneme secretion assays

Microneme secretion assays were conducted as previously described [107]. Freshly egressed parasites were collected, filtered through an 8 µm nuclepore membrane and resuspended in invasion media (20 mM HEPES in DMEM-HG) at 8x10^8^ parasites/mL and stored on ice. 50 µL of parasite suspension was aliquoted for a whole cell control and lysed using CelLytic M [CAT:C2978] with 1 µL DNase I [CAT: EN0521] and 1µL RNase A [CAT:19101]. 100 µL of the parasite suspension was added to 100 µL of invasion media, or invasion media containing 2X inducers (either 2 µM ionomycin or equal volume of DMSO). Suspensions were incubated either for 30 min at 37°C for constitutive secretion or for 3 min at 37°C with inducers or controls for induced secretion. After the incubation the suspension was immediately placed in ice and centrifuged at 1,000 x g at 4°C for 5 minutes. 180 µL of supernatant was placed in a fresh tube and centrifuged again to remove any remaining parasites and 150 µL of supernatant was placed in a fresh tube. A portion of the secretions was boiled with 4x Laemmli for 5 min. Western blots were conducted as described above. Blots were stained with MIC2 and GRA1 to compare MIC2 secretion to the constitutively secreted GRA1 [107].

#### Microneme and rhoptry maturation assay

Microneme maturation assays were conducted as previously described with some modifications [107]. Freshly egressed parasites were collected from wildtype and mutant strains either preincubated with ATc or without ATc. Parasites were filtered through a 5 µm nuclepore membrane, and lysed using CelLytic M. Protein concentrations were measured using BCA and evenly loaded into a 10% SDS-PAGE gel for subsequent western blotting.

Western blots and intracellular immunofluorescence assays were performed in *iΔTgPDIA3* and *TatiΔku80* parasites. Antibodies staining for different secreted proteins (including: mouse-Ty 1:200 (from Drew Ethridge) and rat αHA at 1:50 (Roche) mouse αROP1 at 1:1000 (from BEI resources), mouse αROP7 at 1:2000 (from Peter Bradley), rabbit αMIC2 at 1:1000 (from Vern Carruthers), and mouse αMIC3 at 1:1000 (from BEI resources) were utilized to probe the localization of different rhoptry and microneme secreted proteins in knocked-down parasites (incubated with ATc for 48 hrs) compared to control parasites (0 hrs ATc and *TatiΔku80* parental) [107].

#### Fluorometry measurements of calcium in tachyzoites

For measuring SERCA activity, freshly lysed extracellular tachyzoites were filtered through an 8 µm nuclepore membrane, washed with Buffer A with glucose (BAG) (116 mM NaCl, 5.4 mM KCl, 0.8 mM MgSO_4_, 5.5 mM d-glucose, and 50 mM HEPES, pH 7.4) and resuspended at a concentration of 1x10^9^ parasites/mL in a loading buffer with 20 µM MagFluo-4 AM [CAT: 20401], 0.02% pluronic F127 [CAT: 20052], and 100 µg/mL BSA, in HEPES-buffered saline (HBS) (135 mM NaCl, 5.9 mM KCl, 11.6 mM HEPES, 1.5 mM CaCl_2_, 11.5 mM glucose, 1.2 mM MgCl_2_, pH 7.3) [48, 66]. Parasites were incubated with the indicator for 1 hour with shaking at RT and protected from light. After loading parasites were washed twice with cytosol like media (CLM) (1 mM EGTA, 20 mM PIPES, 20 mM NaCl, 140 mM KCl, pH 7.0) and permeabilized with 100 µM digitonin for 5 minutes. Parasites were washed twice and resuspended in CLM at 1x10^9^ parasites/mL and stored on ice. Fluorescence measurements of 2x10^7^ parasites were taken using a Hitachi F-7000 Fluorescence Spectrophotometer using MagFluo-4 conditions for excitation (495) and emission (528). 220 nM free calcium, added as 375 µM CaCl_2_ (maxchelator for calculations: https://somapp.ucdmc.ucdavis.edu/pharmacology/bers/maxchelator/), 125 µM MgATP, a SERCA substrate, and 1 µM thapsigargin, a SERCA inhibitor, were used to characterize changes in ER calcium sequestration and SERCA function in the absence of TgPDIA3.

For measurement of GECI expressing parasites [65], freshly egressed GCaMP6f-mScarlet mutants were utilized and for mitochondrial calcium measurements force-lysed intracellular GCaMP6f-SOD2 mutants were utilized. For all GECI experiments parasites were collected, washed with BAG, resuspended at 1x10^9^ parasites/mL and kept on ice. Fluorescence measurements were done using a Hitachi F-7000 Fluorescence Spectrophotometer using GCaMP6f conditions for excitation (485) and emission (509) and excitation (569) and emission (594) for mScarlet [65].

Final concentrations of reagents used during fluorometry experiments were as follows: 40 µM GPN, 1 µM thapsigargin, 100 µM zaprinast, 125 µM MgATP, 1 µM ionomycin, and 1.8 mM calcium (for non-permeabilized cells).

#### Ultrastructure expansion microscopy

Expansion microscopy was conducted with intracellular parasites as previously described [114]. In brief, parasites grown on an HFF monolayer on coverslips were fixed 20 hpi, with 4% Paraformaldehyde (PFA) at 37°C for 20 min followed by protein crosslinking prevention with formaldehyde and acrylamide. After 24 hours cells were transferred to sodium acrylate gels (19% sodium acrylate, 10% acrylamide, 0.1% N,N’-methylenbisacrylamide, in PBS pH 7.4, TEMED and APS) and gels were transferred to a denaturing buffer (200 mM SDS, 200 mM NaCl, 50 mM Tris in water, pH9.0) and denatured at 90°C for 90 minutes. Gels were then expanded through three 30-min incubations in water, then shrunken with PBS pH7.4 before blocking with 3% BSA in PBS-T. Samples were incubated overnight with primary antibody in 3% BSA-PBS (rabbit anti- MIC2 at 1:200 (gift from Vern Carruthers), mouse anti-TgPDI1 1:300 (made in-house), and guinea pig anti-SERCA 1:100 (made in-house)). After three washes in PBS-T, the gels were incubated with secondary antibodies and NHS-ester 405 (ThermoFisher) in PBS for 2.5 hours, followed by three additional 30-minute rounds of expansion in water. For imaging, gels were transferred to CellVis dishes [CAT: NC1129240] freshly coated with Poly-D-Lysine to minimize drift or stored at 4°C in 0.2% w/v propyl gallate for up to 48 hours. Imaging was performed on a Zeiss LSM 980 using Airyscan 2, and 3D renderings were generated with IMARIS software.

#### Protein Identification with LC-MS/MS

Samples were submitted to the Proteomics and Metabolomics Facility, Nebraska Center for Biotechnology, University of Nebraska-Lincoln and analyze for mass spectrometry (MS) as previously described [115]. Briefly, an aliquot of 37.5 µL of sample was added to 12.5 µL of 4X reducing NuPAGE LDS gel (Thermo Fisher Scientific, Waltham, MA) sample buffer at 5 mM dithiothreitol (DTT) and incubated at 95°C for 10 min. The samples were loaded and run on a Bolt 12% Bis-Tris-Plus gel (Thermo Fisher Scientific) in MES SDS running buffer to clean them and concentrate the proteins into the top of the gel. The gel was then fixed in methanol:acetic acid:water (40:10:50), and stained with Colloidal Coomassie blue G-250. The gel containing proteins was excised and destained in 50% acetonitrile (ACN), 50 mM ammonium bicarbonate. The proteins were reduced in 100 mM ammonium bicarbonate with DTT at 10 mM. The reducing buffer was removed, and proteins were alkylated with iodoacetamide at 10 mM. Proteins were digested with 250 ng of trypsin overnight at 37°C. Peptides were extracted from the gel pieces, dried down, and re-dissolved in 5% acetonitrile, 0.2% formic acid. Each digest was run by nanoLC-MS/MS using a 2 h gradient on a Waters CSH 0.075 mm x 250 mm C18 column (Waters Corp, Milford, MA) feeding into a Thermo Orbitrap Eclipse mass spectrometer.

All MS/MS samples were analyzed using Mascot (Matrix Science, London, UK; version 2.7). Mascot was set up to search the cRAP_20150130.fasta (125 entries); and ToxoDB- 59_TgondiiGT1_AnnotatedProteins_20221003 (8,460 sequences) assuming the digestion enzyme trypsin. Mascot was searched with a fragment ion mass tolerance of 0.060 Da and a parent ion tolerance of 15.0 PPM. Carbamidomethyl of cysteine was set as a fixed modification. Deamidated of asparagine and glutamine, oxidation of methionine was specified in Mascot as variable modifications. Scaffold (version Scaffold_5.2.2; Proteome Software Inc., Portland, OR) was used to validate LC-MS/MS-based peptide and protein identifications.

Hits were identified with 99% protein threshold, 95% peptide threshold, and a minimum peptide number of 2. Statistical analysis was conducted using Fisher’s exact test with Benjamini-Hochberg multiple test corrections to identify significantly enriched proteins.

The mass spectrometry proteomics data have been deposited to the ProteomeXchange Consortium via the PRIDE [116] partner repository with the dataset identifiers SERCA-HA DVSF HA-IP: PXD063659, streptavadin-IP: PXD063619, and DVSF anti-TgPDIA3-IP: PXD063615.

### Statistical Analysis

Experimental data were expressed as the mean with standard error (SEM) from at least 3 biological replicates unless otherwise indicated. Statistical analyses were completed using GraphPad PRISM. Student’s t-test, one-way ANOVA, two-way ANOVA, and linear regression were used when necessary and are indicated in figure legends. A p-value of <0.05 was considered statistically significant. For all LC-MS/MS analyses, in Scaffold, protein thresholds were set to 95%, a minimum peptide number was set to 2, and peptide threshold was set to 99%. Fisher’s exact test (with Benjamini-Hochberg multiple testing corrections) was used to calculate statistical significance. Minimum values were set to 0.0001 to avoid infinity.

## Acknowledgments

We thank the Proteomics & Metabolomics Facility (RRID:SCR_021314), Nebraska Center for Biotechnology at the University of Nebraska-Lincoln for the LC-MS/MS analysis. The facility and instrumentation are supported by the Nebraska Research Initiative. We thank the CTEGD FACS sorting and biomedical microscopy cores for the use of their facilities and equipment. Julie Nelson and Muthugapatti K Kandasamy provided training and support. We thank Andrea Hortua Triana, Karla Marquez Nogueras, and Abigail Calixto for training on the use of *T. gondii* protocols. Mayara Bertolini, Baihetiya Baierna, and Melissa Sleda for advice on protein purification, subcellular fractionation and streptavidin immunoprecipitation, and IFAs, respectively. Zhuhong Li and Andrea Hortua Triana provided training and advice in calcium measurements. Anna Gioseffi for the training on the use of the scaffold software. Juan Camilo Arenas Garcia for help with the phylogenetic analyses. Andrea Hortua Triana directed the generation of the αTgPDIA3 and αGRA1 antibodies. Peter Bradley, Vern Carruthers, Drew Ethridge, and BEI resources provided various antibodies. We thank Diego Huet for the IP protocol used for the TgPDIA3 DVSF IP. The *FBXO14-TID* cell line was a gift from Diego Huet’s lab. Vasant Muralidharan lab and David Cobb allowed the use of their reagents and DVSF cross-linking protocol. Vasant Muralidharan lab and Grace Vick assisted with the ultrastructure expansion microscopy protocol and training. Drew Etheridge lab and Justin Wiedeman for the elution of biotinylated proteins from streptavidin beads. This work was funded by NIH grants AI154931 and 174600 to SNJM. KEM was partially supported by a pre-doc fellowship funded by AIT32060546.

## References

1. Barylyuk K, Koreny L, Ke H, Butterworth S, Crook OM, Lassadi I, et al. A Comprehensive Subcellular Atlas of the Toxoplasma Proteome via hyperLOPIT Provides Spatial Context for Protein Functions. Cell Host Microbe. 2020;28(5):752–66 e9. Epub 20201013. doi: 10.1016/j.chom.2020.09.011. PubMed PMID: 33053376; PubMed Central PMCID: PMCPMC7670262.

2. Hill D, Dubey JP. Toxoplasma gondii: transmission, diagnosis and prevention. Clin Microbiol Infect. 2002;8(10):634–40. doi: 10.1046/j.1469-0691.2002.00485.x. PubMed PMID: 12390281.

3. Jones JL, Parise ME, Fiore AE. Neglected parasitic infections in the United States: toxoplasmosis. Am J Trop Med Hyg. 2014;90(5):794–9. doi: 10.4269/ajtmh.13-0722. PubMed PMID: 24808246; PubMed Central PMCID: PMCPMC4015566.

4. Weiss LM, Kim K. The development and biology of bradyzoites of Toxoplasma gondii. Front Biosci. 2000;5:D391-405. Epub 20000401. doi: 10.2741/weiss. PubMed PMID: 10762601; PubMed Central PMCID: PMCPMC3109641.

5. Luft BJ, Remington JS. Toxoplasmic encephalitis in AIDS. Clin Infect Dis. 1992;15(2):211–22. doi: 10.1093/clinids/15.2.211. PubMed PMID: 1520757.

6. Hampton MM. Congenital Toxoplasmosis: A Review. Neonatal Netw. 2015;34(5):274-8. doi: 10.1891/0730-0832.34.5.274. PubMed PMID: WOS:000440840400003.

7. Black MW, Boothroyd JC. Lytic cycle of Toxoplasma gondii. Microbiol Mol Biol Rev. 2000;64(3):607–23. doi: 10.1128/MMBR.64.3.607-623.2000. PubMed PMID: 10974128; PubMed Central PMCID: PMCPMC99006.

8. Blader IJ, Coleman BI, Chen CT, Gubbels MJ. Lytic Cycle of Toxoplasma gondii: 15 Years Later. Annu Rev Microbiol. 2015;69:463–85. Epub 20150828. doi: 10.1146/annurev-micro-091014-104100. PubMed PMID: 26332089; PubMed Central PMCID: PMCPMC4659696.

9. Carruthers VB, Giddings OK, Sibley LD. Secretion of micronemal proteins is associated with toxoplasma invasion of host cells. Cell Microbiol. 1999;1(3):225–35. doi: 10.1046/j.1462-5822.1999.00023.x. PubMed PMID: 11207555.

10. Boothroyd JC, Dubremetz JF. Kiss and spit: the dual roles of Toxoplasma rhoptries. Nat Rev Microbiol. 2008;6(1):79–88. doi: 10.1038/nrmicro1800. PubMed PMID: 18059289.

11. Carruthers VB, Sibley LD. Mobilization of intracellular calcium stimulates microneme discharge in Toxoplasma gondii. Mol Microbiol. 1999;31(2):421–8. doi: 10.1046/j.1365-2958.1999.01174.x. PubMed PMID: 10027960.

12. Clapham DE. Calcium signaling. Cell. 2007;131(6):1047-58. Epub 2007/12/18. doi: 10.1016/j.cell.2007.11.028. PubMed PMID: 18083096.

13. Bootman MD, Bultynck G. Fundamentals of Cellular Calcium Signaling: A Primer. Cold Spring Harb Perspect Biol. 2020;12(1). Epub 20200102. doi: 10.1101/cshperspect.a038802. PubMed PMID: 31427372; PubMed Central PMCID: PMCPMC6942118.

14. Bootman MD, Petersen OH, Verkhratsky A. The endoplasmic reticulum is a focal point for co-ordination of cellular activity. Cell Calcium. 2002;32(5-6):231–4. doi: 10.1016/S0143416002002002. PubMed PMID: WOS:000180877600001.

15. Viotti C. ER to Golgi-Dependent Protein Secretion: The Conventional Pathway. Methods Mol Biol. 2016;1459:3–29. doi: 10.1007/978-1-4939-3804-9_1. PubMed PMID: WOS:000687281000002.

16. Ni M, Lee AS. ER chaperones in mammalian development and human diseases. FEBS Lett. 2007;581(19):3641–51. Epub 20070425. doi: 10.1016/j.febslet.2007.04.045. PubMed PMID: 17481612; PubMed Central PMCID: PMCPMC2040386.

17. Meldolesi J, Pozzan T. The endoplasmic reticulum Ca2+ store: a view from the lumen. Trends Biochem Sci. 1998;23(1):10–4. doi: 10.1016/s0968-0004(97)01143-2. PubMed PMID: 9478128.

18. Michalak M, Robert Parker JM, Opas M. Ca2+ signaling and calcium binding chaperones of the endoplasmic reticulum. Cell Calcium. 2002;32(5-6):269–78. doi: 10.1016/s0143416002001884. PubMed PMID: 12543089.

19. Hwang C, Sinskey AJ, Lodish HF. Oxidized redox state of glutathione in the endoplasmic reticulum. Science. 1992;257(5076):1496-502. doi: 10.1126/science.1523409. PubMed PMID: 1523409.

20. Ali Khan H, Mutus B. Protein disulfide isomerase a multifunctional protein with multiple physiological roles. Front Chem. 2014;2:70. Epub 20140826. doi: 10.3389/fchem.2014.00070. PubMed PMID: 25207270; PubMed Central PMCID: PMCPMC4144422.

21. Galligan JJ, Petersen DR. The human protein disulfide isomerase gene family. Hum Genomics. 2012;6(1):6. Epub 20120705. doi: 10.1186/1479-7364-6-6. PubMed PMID: 23245351; PubMed Central PMCID: PMCPMC3500226.

22. Wang L, Wang X, Wang CC. Protein disulfide-isomerase, a folding catalyst and a redox-regulated chaperone. Free Radic Biol Med. 2015;83:305–13. Epub 20150217. doi: 10.1016/j.freeradbiomed.2015.02.007. PubMed PMID: 25697778.

23. Chen X, Shi C, He M, Xiong S, Xia X. Endoplasmic reticulum stress: molecular mechanism and therapeutic targets. Signal Transduct Target Ther. 2023;8(1):352. Epub 20230915. doi: 10.1038/s41392-023-01570-w. PubMed PMID: 37709773; PubMed Central PMCID: PMCPMC10502142.

24. Wang L, Zhou J, Wang L, Wang CC, Essex DW. The b’ domain of protein disulfide isomerase cooperates with the a and a’ domains to functionally interact with platelets. J Thromb Haemost. 2019;17(2):371–82. Epub 20190203. doi: 10.1111/jth.14366. PubMed PMID: 30566278; PubMed Central PMCID: PMCPMC6368866.

25. Kozlov G, Maattanen P, Schrag JD, Pollock S, Cygler M, Nagar B, et al. Crystal structure of the bb’ domains of the protein disulfide isomerase ERp57. Structure. 2006;14(8):1331–9. doi: 10.1016/j.str.2006.06.019. PubMed PMID: 16905107.

26. Maattanen P, Kozlov G, Gehring K, Thomas DY. ERp57 and PDI: multifunctional protein disulfide isomerases with similar domain architectures but differing substrate-partner associations. Biochem Cell Biol. 2006;84(6):881–9. doi: 10.1139/o06-186. PubMed PMID: 17215875.

27. Bastos-Aristizabal S, Kozlov G, Gehring K. Structure of the substrate-binding b’ domain of the Protein Disulfide Isomerase-Like protein of the Testis. Sci Rep. 2014;4:4464. Epub 20140325. doi: 10.1038/srep04464. PubMed PMID: 24662985; PubMed Central PMCID: PMCPMC4894388.

28. Ushioda R, Miyamoto A, Inoue M, Watanabe S, Okumura M, Maegawa KI, et al. Redox- assisted regulation of Ca2+ homeostasis in the endoplasmic reticulum by disulfide reductase ERdj5. Proc Natl Acad Sci U S A. 2016;113(41):E6055-E63. Epub 20160930. doi: 10.1073/pnas.1605818113. PubMed PMID: 27694578; PubMed Central PMCID: PMCPMC5068290.

29. Li G, Mongillo M, Chin KT, Harding H, Ron D, Marks AR, et al. Role of ERO1-alpha- mediated stimulation of inositol 1,4,5-triphosphate receptor activity in endoplasmic reticulum stress-induced apoptosis. J Cell Biol. 2009;186(6):783-92. Epub 20090914. doi: 10.1083/jcb.200904060. PubMed PMID: 19752026; PubMed Central PMCID: PMCPMC2753154.

30. Higo T, Hattori M, Nakamura T, Natsume T, Michikawa T, Mikoshiba K. Subtype-specific and ER lumenal environment-dependent regulation of inositol 1,4,5-trisphosphate receptor type 1 by ERp44. Cell. 2005;120(1):85-98. doi: 10.1016/j.cell.2004.11.048. PubMed PMID: 15652484.

31. Li Y, Camacho P. Ca-dependent redox modulation of SERCA 2b by ERp57. J Cell Biol. 2004;164(1):35–46. doi: DOI 10.1083/jcb.200307010. PubMed PMID: WOS:000188065500005.

32. Fani G, La Torre CE, Cascella R, Cecchi C, Vendruscolo M, Chiti F. Misfolded protein oligomers induce an increase of intracellular Ca(2+) causing an escalation of reactive oxidative species. Cell Mol Life Sci. 2022;79(9):500. Epub 20220827. doi: 10.1007/s00018-022-04513-w. PubMed PMID: 36030306; PubMed Central PMCID: PMCPMC9420098.

33. Makio T, Chen J, Simmen T. ER stress as a sentinel mechanism for ER Ca(2+) homeostasis. Cell Calcium. 2024;124:102961. Epub 20241018. doi: 10.1016/j.ceca.2024.102961. PubMed PMID: 39471738.

34. Weids AJ, Ibstedt S, Tamas MJ, Grant CM. Distinct stress conditions result in aggregation of proteins with similar properties. Sci Rep. 2016;6:24554. Epub 20160418. doi: 10.1038/srep24554. PubMed PMID: 27086931; PubMed Central PMCID: PMCPMC4834537.

35. Shenton D, Grant CM. Protein S-thiolation targets glycolysis and protein synthesis in response to oxidative stress in the yeast Saccharomyces cerevisiae. Biochem J. 2003;374(Pt 2):513–9. doi: 10.1042/BJ20030414. PubMed PMID: 12755685; PubMed Central PMCID: PMCPMC1223596.

36. Raturi A, Gutierrez T, Ortiz-Sandoval C, Ruangkittisakul A, Herrera-Cruz MS, Rockley JP, et al. TMX1 determines cancer cell metabolism as a thiol-based modulator of ER- mitochondria Ca2+ flux. J Cell Biol. 2016;214(4):433–44. Epub 20160808. doi: 10.1083/jcb.201512077. PubMed PMID: 27502484; PubMed Central PMCID: PMCPMC4987292.

37. Lynes EM, Raturi A, Shenkman M, Ortiz Sandoval C, Yap MC, Wu J, et al. Palmitoylation is the switch that assigns calnexin to quality control or ER Ca2+ signaling. J Cell Sci. 2013;126(Pt 17):3893–903. Epub 20130710. doi: 10.1242/jcs.125856. PubMed PMID: 23843619.

38. Cao SS, Kaufman RJ. Endoplasmic reticulum stress and oxidative stress in cell fate decision and human disease. Antioxid Redox Signal. 2014;21(3):396–413. Epub 20140612. doi: 10.1089/ars.2014.5851. PubMed PMID: 24702237; PubMed Central PMCID: PMCPMC4076992.

39. Soares Moretti AI, Martins Laurindo FR. Protein disulfide isomerases: Redox connections in and out of the endoplasmic reticulum. Arch Biochem Biophys. 2017;617:106–19. Epub 20161124. doi: 10.1016/j.abb.2016.11.007. PubMed PMID: 27889386.

40. Avezov E, Konno T, Zyryanova A, Chen W, Laine R, Crespillo-Casado A, et al. Retarded PDI diffusion and a reductive shift in poise of the calcium depleted endoplasmic reticulum. Bmc Biol. 2015;13. doi: ARTN 210.1186/s12915-014-0112-2. PubMed PMID: WOS:000349191000001.

41. Chichiarelli S, Altieri F, Paglia G, Rubini E, Minacori M, Eufemi M. ERp57/PDIA3: new insight. Cell Mol Biol Lett. 2022;27(1):12. Epub 20220202. doi: 10.1186/s11658-022-00315-x. PubMed PMID: 35109791; PubMed Central PMCID: PMCPMC8809632.

42. Sidik SM, Huet D, Ganesan SM, Huynh MH, Wang T, Nasamu AS, et al. A Genome-wide CRISPR Screen in Toxoplasma Identifies Essential Apicomplexan Genes. Cell. 2016;166(6):1423–35 e12. Epub 20160902. doi: 10.1016/j.cell.2016.08.019. PubMed PMID: 27594426; PubMed Central PMCID: PMCPMC5017925.

43. Pinto RD, Moreira AR, Pereira PJ, dos Santos NM. Two thioredoxin-superfamily members from sea bass (Dicentrarchus labrax, L.): characterization of PDI (PDIA1) and ERp57 (PDIA3). Fish Shellfish Immunol. 2013;35(4):1163-75. Epub 20130721. doi: 10.1016/j.fsi.2013.07.024. PubMed PMID: 23880452.

44. Jumper J, Evans R, Pritzel A, Green T, Figurnov M, Ronneberger O, et al. Highly accurate protein structure prediction with AlphaFold. Nature. 2021;596(7873):583-9. Epub 20210715. doi: 10.1038/s41586-021-03819-2. PubMed PMID: 34265844; PubMed Central PMCID: PMCPMC8371605.

45. Varadi M, Bertoni D, Magana P, Paramval U, Pidruchna I, Radhakrishnan M, et al. AlphaFold Protein Structure Database in 2024: providing structure coverage for over 214 million protein sequences. Nucleic Acids Res. 2024;52(D1):D368–D75. doi: 10.1093/nar/gkad1011. PubMed PMID: 37933859; PubMed Central PMCID: PMCPMC10767828.

46. Muller IK, Winter C, Thomas C, Spaapen RM, Trowitzsch S, Tampe R. Structure of an MHC I-tapasin-ERp57 editing complex defines chaperone promiscuity. Nat Commun. 2022;13(1):5383. Epub 20220914. doi: 10.1038/s41467-022-32841-9. PubMed PMID: 36104323; PubMed Central PMCID: PMCPMC9474470.

47. Dong G, Wearsch PA, Peaper DR, Cresswell P, Reinisch KM. Insights into MHC class I peptide loading from the structure of the tapasin-ERp57 thiol oxidoreductase heterodimer. Immunity. 2009;30(1):21–32. doi: 10.1016/j.immuni.2008.10.018. PubMed PMID: 19119025; PubMed Central PMCID: PMCPMC2650231.

48. Li ZH, Asady B, Chang L, Triana MAH, Li C, Coppens I, et al. Calcium transfer from the ER to other organelles for optimal signaling in Toxoplasma gondii. bioRxiv. 2024. Epub 20241205. doi: 10.1101/2024.08.15.608087. PubMed PMID: 39185237; PubMed Central PMCID: PMCPMC11343207.

49. Sheiner L, Demerly JL, Poulsen N, Beatty WL, Lucas O, Behnke MS, et al. A systematic screen to discover and analyze apicoplast proteins identifies a conserved and essential protein import factor. PLoS Pathog. 2011;7(12):e1002392. Epub 20111201. doi: 10.1371/journal.ppat.1002392. PubMed PMID: 22144892; PubMed Central PMCID: PMCPMC3228799.

50. Kafsack BF, Beckers C, Carruthers VB. Synchronous invasion of host cells by Toxoplasma gondii. Mol Biochem Parasitol. 2004;136(2):309–11. doi: 10.1016/j.molbiopara.2004.04.004. PubMed PMID: 15478810.

51. Chasen NM, Asady B, Lemgruber L, Vommaro RC, Kissinger JC, Coppens I, et al. A Glycosylphosphatidylinositol-Anchored Carbonic Anhydrase-Related Protein of Toxoplasma gondii Is Important for Rhoptry Biogenesis and Virulence. mSphere. 2017;2(3). Epub 20170517. doi: 10.1128/mSphere.00027-17. PubMed PMID: 28529974; PubMed Central PMCID: PMCPMC5437132.

52. Dahiya V, Buchner J. Functional principles and regulation of molecular chaperones. Adv Protein Chem Struct Biol. 2019;114:1–60. Epub 20181201. doi: 10.1016/bs.apcsb.2018.10.001. PubMed PMID: 30635079.

53. Mares RE, Melendez-Lopez SG, Ramos MA. Acid-denatured Green Fluorescent Protein (GFP) as model substrate to study the chaperone activity of protein disulfide isomerase. Int J Mol Sci. 2011;12(7):4625–36. Epub 20110718. doi: 10.3390/ijms12074625. PubMed PMID: 21845100; PubMed Central PMCID: PMCPMC3155373.

54. Allan KM, Loberg MA, Chepngeno J, Hurtig JE, Tripathi S, Kang MG, et al. Trapping redox partnerships in oxidant-sensitive proteins with a small, thiol-reactive cross-linker. Free Radic Biol Med. 2016;101:356–66. Epub 20161102. doi: 10.1016/j.freeradbiomed.2016.10.506. PubMed PMID: 27816612; PubMed Central PMCID: PMCPMC5154803.

55. Araki K, Ushioda R, Kusano H, Tanaka R, Hatta T, Fukui K, et al. A crosslinker-based identification of redox relay targets. Anal Biochem. 2017;520:22–6. Epub 20161231. doi: 10.1016/j.ab.2016.12.025. PubMed PMID: 28048978.

56. Naticchia MR, Brown HA, Garcia FJ, Lamade AM, Justice SL, Herrin RP, et al. Bifunctional electrophiles cross-link thioredoxins with redox relay partners in cells. Chem Res Toxicol. 2013;26(3):490–7. Epub 20130304. doi: 10.1021/tx4000123. PubMed PMID: 23414292; PubMed Central PMCID: PMCPMC3637938.

57. Wang L, Li SJ, Sidhu A, Zhu L, Liang Y, Freedman RB, et al. Reconstitution of human Ero1- Lalpha/protein-disulfide isomerase oxidative folding pathway in vitro. Position-dependent differences in role between the a and a’ domains of protein-disulfide isomerase. J Biol Chem. 2009;284(1):199–206. Epub 20081111. doi: 10.1074/jbc.M806645200. PubMed PMID: 19001419.

58. Alanen HI, Williamson RA, Howard MJ, Hatahet FS, Salo KE, Kauppila A, et al. ERp27, a new non-catalytic endoplasmic reticulum-located human protein disulfide isomerase family member, interacts with ERp57. J Biol Chem. 2006;281(44):33727–38. Epub 20060828. doi: 10.1074/jbc.M604314200. PubMed PMID: 16940051.

59. Sugiura Y, Araki K, Iemura S, Natsume T, Hoseki J, Nagata K. Novel thioredoxin-related transmembrane protein TMX4 has reductase activity. J Biol Chem. 2010;285(10):7135–42. Epub 20100107. doi: 10.1074/jbc.M109.082545. PubMed PMID: 20056998; PubMed Central PMCID: PMCPMC2844163.

60. Mandalasi MN, Gas-Pascual E, Baptista CG, Deng B, van der Wel H, Kruijtzer JAW, et al. Oxygen-dependent regulation of F-box proteins in Toxoplasma gondii is mediated by Skp1 glycosylation. J Biol Chem. 2024;300(11):107801. Epub 20240921. doi: 10.1016/j.jbc.2024.107801. PubMed PMID: 39307307; PubMed Central PMCID: PMCPMC11570480.

61. Sheiner L, Soldati-Favre D. Protein trafficking inside Toxoplasma gondii. Traffic. 2008;9(5):636–46. Epub 20080304. doi: 10.1111/j.1600-0854.2008.00713.x. PubMed PMID: 18331382.

62. Lovett JL, Marchesini N, Moreno SN, Sibley LD. Toxoplasma gondii microneme secretion involves intracellular Ca(2+) release from inositol 1,4,5-triphosphate (IP(3))/ryanodine-sensitive stores. J Biol Chem. 2002;277(29):25870-6. Epub 20020513. doi: 10.1074/jbc.M202553200. PubMed PMID: 12011085.

63. Song G, Springer TA. Structures of the Toxoplasma gliding motility adhesin. Proc Natl Acad Sci U S A. 2014;111(13):4862–7. Epub 20140317. doi: 10.1073/pnas.1403059111. PubMed PMID: 24639528; PubMed Central PMCID: PMCPMC3977308.

64. John LM, Lechleiter JD, Camacho P. Differential modulation of SERCA2 isoforms by calreticulin. J Cell Biol. 1998;142(4):963–73. doi: 10.1083/jcb.142.4.963. PubMed PMID: 9722609; PubMed Central PMCID: PMCPMC2132884.

65. Vella SA, Calixto A, Asady B, Li ZH, Moreno SNJ. Genetic Indicators for Calcium Signaling Studies in Toxoplasma gondii. Methods Mol Biol. 2020;2071:187–207. doi: 10.1007/978-1-4939-9857-9_11. PubMed PMID: 31758454; PubMed Central PMCID: PMCPMC7294879.

66. Rossi AM, Taylor CW. Reliable measurement of free Ca(2+) concentrations in the ER lumen using Mag-Fluo-4. Cell Calcium. 2020;87:102188. Epub 20200306. doi: 10.1016/j.ceca.2020.102188. PubMed PMID: 32179239; PubMed Central PMCID: PMCPMC7181174.

67. Calixto A, Moen K, Moreno SN. The contribution of the Golgi and the endoplasmic reticulum to calcium and pH homeostasis in Toxoplasma gondii. J Biol Chem. 2025:108372. Epub 20250303. doi: 10.1016/j.jbc.2025.108372. PubMed PMID: 40043955.

68. Marino M, Stoilova T, Giorgi C, Bachi A, Cattaneo A, Auricchio A, et al. SEPN1, an endoplasmic reticulum-localized selenoprotein linked to skeletal muscle pathology, counteracts hyperoxidation by means of redox-regulating SERCA2 pump activity. Hum Mol Genet. 2015;24(7):1843–55. Epub 20141201. doi: 10.1093/hmg/ddu602. PubMed PMID: 25452428.

69. Ramakrishnan C, Maier S, Walker RA, Rehrauer H, Joekel DE, Winiger RR, et al. An experimental genetically attenuated live vaccine to prevent transmission of Toxoplasma gondii by cats. Sci Rep. 2019;9(1):1474. Epub 20190206. doi: 10.1038/s41598-018-37671-8. PubMed PMID: 30728393; PubMed Central PMCID: PMCPMC6365665.

70. Chen TW, Wardill TJ, Sun Y, Pulver SR, Renninger SL, Baohan A, et al. Ultrasensitive fluorescent proteins for imaging neuronal activity. Nature. 2013;499(7458):295-300. doi: 10.1038/nature12354. PubMed PMID: 23868258; PubMed Central PMCID: PMCPMC3777791.

71. Pino P, Foth BJ, Kwok LY, Sheiner L, Schepers R, Soldati T, et al. Dual targeting of antioxidant and metabolic enzymes to the mitochondrion and the apicoplast of Toxoplasma gondii. PLoS Pathog. 2007;3(8):e115. doi: 10.1371/journal.ppat.0030115. PubMed PMID: 17784785; PubMed Central PMCID: PMCPMC1959373.

72. Haller T, Dietl P, Deetjen P, Volkl H. The lysosomal compartment as intracellular calcium store in MDCK cells: a possible involvement in InsP3-mediated Ca2+ release. Cell Calcium. 1996;19(2):157–65. doi: 10.1016/s0143-4160(96)90084-6. PubMed PMID: 8689673.

73. Miranda K, Pace DA, Cintron R, Rodrigues JC, Fang J, Smith A, et al. Characterization of a novel organelle in Toxoplasma gondii with similar composition and function to the plant vacuole. Mol Microbiol. 2010;76(6):1358–75. Epub 20100414. doi: 10.1111/j.1365-2958.2010.07165.x. PubMed PMID: 20398214; PubMed Central PMCID: PMCPMC2907454.

74. Stasic AJ, Moreno SNJ, Carruthers VB, Dou Z. The Toxoplasma plant-like vacuolar compartment (PLVAC). J Eukaryot Microbiol. 2022;69(6):e12951. Epub 20221027. doi: 10.1111/jeu.12951. PubMed PMID: 36218001; PubMed Central PMCID: PMCPMC10576567.

75. Ostergaard H, Henriksen A, Hansen FG, Winther JR. Shedding light on disulfide bond formation: engineering a redox switch in green fluorescent protein. EMBO J. 2001;20(21):5853–62. doi: 10.1093/emboj/20.21.5853. PubMed PMID: 11689426; PubMed Central PMCID: PMCPMC125700.

76. Cai H, Wang CC, Tsou CL. Chaperone-like activity of protein disulfide isomerase in the refolding of a protein with no disulfide bonds. J Biol Chem. 1994;269(40):24550–2. PubMed PMID: 7929125.

77. Cobb DW, Kudyba HM, Villegas A, Hoopmann MR, Baptista RP, Bruton B, et al. A redox-active crosslinker reveals an essential and inhibitable oxidative folding network in the endoplasmic reticulum of malaria parasites. PLoS Pathog. 2021;17(2):e1009293. Epub 20210203. doi: 10.1371/journal.ppat.1009293. PubMed PMID: 33534803; PubMed Central PMCID: PMCPMC7886143.

78. Zhang L, Niu Y, Zhu L, Fang J, Wang X, Wang L, et al. Different interaction modes for protein-disulfide isomerase (PDI) as an efficient regulator and a specific substrate of endoplasmic reticulum oxidoreductin-1alpha (Ero1alpha). J Biol Chem. 2014;289(45):31188–99. Epub 20140925. doi: 10.1074/jbc.M114.602961. PubMed PMID: 25258311; PubMed Central PMCID: PMCPMC4223321.

79. Ramirez-Flores CJ, Cruz-Miron R, Arroyo R, Mondragon-Castelan ME, Nopal-Guerrero T, Gonzalez-Pozos S, et al. Characterization of metalloproteases and serine proteases of Toxoplasma gondii tachyzoites and their effect on epithelial cells. Parasitol Res. 2019;118(1):289–306. Epub 20181201. doi: 10.1007/s00436-018-6163-5. PubMed PMID: 30506516.

80. Hajagos BE, Turetzky JM, Peng ED, Cheng SJ, Ryan CM, Souda P, et al. Molecular dissection of novel trafficking and processing of the Toxoplasma gondii rhoptry metalloprotease toxolysin-1. Traffic. 2012;13(2):292–304. Epub 20111129. doi: 10.1111/j.1600-0854.2011.01308.x. PubMed PMID: 22035499; PubMed Central PMCID: PMCPMC3375832.

81. Laliberte J, Carruthers VB. Toxoplasma gondii toxolysin 4 is an extensively processed putative metalloproteinase secreted from micronemes. Mol Biochem Parasitol. 2011;177(1):49–56. Epub 20110126. doi: 10.1016/j.molbiopara.2011.01.009. PubMed PMID: 21277910; PubMed Central PMCID: PMCPMC3057334.

82. Escotte-Binet S, Huguenin A, Aubert D, Martin AP, Kaltenbach M, Florent I, et al. Metallopeptidases of Toxoplasma gondii: in silico identification and gene expression. Parasite. 2018;25:26. Epub 20180508. doi: 10.1051/parasite/2018025. PubMed PMID: 29737275; PubMed Central PMCID: PMCPMC5939537.

83. Wolf DH, Stolz A. The Cdc48 machine in endoplasmic reticulum associated protein degradation. Biochim Biophys Acta. 2012;1823(1):117–24. Epub 20110916. doi: 10.1016/j.bbamcr.2011.09.002. PubMed PMID: 21945179.

84. Carruthers VB. Armed and dangerous: Toxoplasma gondii uses an arsenal of secretory proteins to infect host cells. Parasitol Int. 1999;48(1):1–10. doi: 10.1016/s1383-5769(98)00042-7. PubMed PMID: 11269320.

85. Venugopal K, Marion S. Secretory organelle trafficking in Toxoplasma gondii: A long story for a short travel. Int J Med Microbiol. 2018;308(7):751–60. Epub 20180721. doi: 10.1016/j.ijmm.2018.07.007. PubMed PMID: 30055977.

86. Wang C, Sun P, Jia Y, Tang X, Liu X, Suo X, et al. Protein disulfide isomerase PDI8 is indispensable for parasite growth and associated with secretory protein processing in Toxoplasma gondii. mBio. 2024;15(9):e0205124. Epub 20240820. doi: 10.1128/mbio.02051-24. PubMed PMID: 39162526; PubMed Central PMCID: PMCPMC11389393.

87. Nagamune K, Beatty WL, Sibley LD. Artemisinin induces calcium-dependent protein secretion in the protozoan parasite. Eukaryot Cell. 2007;6(11):2147–56. doi: 10.1128/Ec.00262-07. PubMed PMID: WOS:000251410200022.

88. Berridge MJ. Calcium microdomains: organization and function. Cell Calcium. 2006;40(5-6):405-12. Epub 20061009. doi: 10.1016/j.ceca.2006.09.002. PubMed PMID: 17030366.

89. Barak P, Parekh AB. Signaling through Ca(2+) Microdomains from Store-Operated CRAC Channels. Cold Spring Harb Perspect Biol. 2020;12(7). Epub 20200701. doi: 10.1101/cshperspect.a035097. PubMed PMID: 31358516; PubMed Central PMCID: PMCPMC7328460.

90. Meyer BA, Doroudgar S. ER Stress-Induced Secretion of Proteins and Their Extracellular Functions in the Heart. Cells. 2020;9(9). Epub 20200910. doi: 10.3390/cells9092066. PubMed PMID: 32927693; PubMed Central PMCID: PMCPMC7563782.

91. Peters LR, Raghavan M. Endoplasmic reticulum calcium depletion impacts chaperone secretion, innate immunity, and phagocytic uptake of cells. J Immunol. 2011;187(2):919–31. Epub 20110613. doi: 10.4049/jimmunol.1100690. PubMed PMID: 21670312; PubMed Central PMCID: PMCPMC3371385.

92. Sharda A, Kim SH, Jasuja R, Gopal S, Flaumenhaft R, Furie BC, et al. Defective PDI release from platelets and endothelial cells impairs thrombus formation in Hermansky-Pudlak syndrome. Blood. 2015;125(10):1633–42. Epub 20150115. doi: 10.1182/blood-2014-08-597419. PubMed PMID: 25593336; PubMed Central PMCID: PMCPMC4351508.

93. Cho J, Furie BC, Coughlin SR, Furie B. A critical role for extracellular protein disulfide isomerase during thrombus formation in mice. J Clin Invest. 2008;118(3):1123–31. doi: 10.1172/JCI34134. PubMed PMID: 18292814; PubMed Central PMCID: PMCPMC2248441.

94. Yavuz U, Alaylioglu M, Sengul B, Karras SN, Gezen-Ak D, Dursun E. Protein disulfide isomerase A3 might be involved in the regulation of 24-dehydrocholesterol reductase via vitamin D equilibrium in primary cortical neurons. In Vitro Cell Dev Biol Anim. 2021;57(7):704–14. Epub 20210802. doi: 10.1007/s11626-021-00602-5. PubMed PMID: 34338991.

95. Nowak JI, Olszewska AM, Krol O, Zmijewski MA. Protein Disulfide Isomerase Family A Member 3 Knockout Abrogate Effects of Vitamin D on Cellular Respiration and Glycolysis in Squamous Cell Carcinoma. Nutrients. 2023;15(21). Epub 20231025. doi: 10.3390/nu15214529. PubMed PMID: 37960182; PubMed Central PMCID: PMCPMC10650882.

96. Meek B, Back JW, Klaren VN, Speijer D, Peek R. Protein disulfide isomerase of Toxoplasma gondii is targeted by mucosal IgA antibodies in humans. FEBS Lett. 2002;522(1-3):104–8. doi: 10.1016/s0014-5793(02)02911-3. PubMed PMID: 12095627.

97. Moncada D, Arenas A, Acosta A, Molina D, Hernandez A, Cardona N, et al. Role of the 52 KDa thioredoxin protein disulfide isomerase of Toxoplasma gondii during infection to human cells. Exp Parasitol. 2016;164:36–42. Epub 20160217. doi: 10.1016/j.exppara.2016.02.005. PubMed PMID: 26896642.

98. Lynes EM, Bui M, Yap MC, Benson MD, Schneider B, Ellgaard L, et al. Palmitoylated TMX and calnexin target to the mitochondria-associated membrane. EMBO J. 2012;31(2):457–70. Epub 20111101. doi: 10.1038/emboj.2011.384. PubMed PMID: 22045338; PubMed Central PMCID: PMCPMC3261551.

99. Borges-Pereira L, Budu A, McKnight CA, Moore CA, Vella SA, Hortua Triana MA, et al. Calcium Signaling throughout the Toxoplasma gondii Lytic Cycle: A STUDY USING GENETICALLY ENCODED CALCIUM INDICATORS. J Biol Chem. 2015;290(45):26914-26. Epub 20150915. doi: 10.1074/jbc.M115.652511. PubMed PMID: 26374900; PubMed Central PMCID: PMCPMC4646405.

100. Lourido S, Moreno SNJ. The calcium signaling toolkit of the Apicomplexan parasites and spp. Cell Calcium. 2015;57(3):186–93. doi: 10.1016/j.ceca.2014.12.010. PubMed PMID: WOS:000351808900008.

101. Hortua Triana MA, Marquez-Nogueras KM, Vella SA, Moreno SNJ. Calcium signaling and the lytic cycle of the Apicomplexan parasite Toxoplasma gondii. Biochim Biophys Acta Mol Cell Res. 2018;1865(11 Pt B):1846-56. Epub 20180810. doi: 10.1016/j.bbamcr.2018.08.004. PubMed PMID: 30992126; PubMed Central PMCID: PMCPMC6477927.

102. Vella SA, Moore CA, Li ZH, Hortua Triana MA, Potapenko E, Moreno SNJ. The role of potassium and host calcium signaling in Toxoplasma gondii egress. Cell Calcium. 2021;94:102337. Epub 20210119. doi: 10.1016/j.ceca.2020.102337. PubMed PMID: 33524795; PubMed Central PMCID: PMCPMC7914212.

103. Alvarez-Jarreta J, Amos B, Aurrecoechea C, Bah S, Barba M, Barreto A, et al. VEuPathDB: the eukaryotic pathogen, vector and host bioinformatics resource center in 2023. Nucleic Acids Res. 2024;52(D1):D808–D16. doi: 10.1093/nar/gkad1003. PubMed PMID: 37953350; PubMed Central PMCID: PMCPMC10767879.

104. Tamura K, Stecher G, Kumar S. MEGA11: Molecular Evolutionary Genetics Analysis Version 11. Mol Biol Evol. 2021;38(7):3022–7. doi: 10.1093/molbev/msab120. PubMed PMID: 33892491; PubMed Central PMCID: PMCPMC8233496.

105. Shen B, Brown K, Long S, Sibley LD. Development of CRISPR/Cas9 for Efficient Genome Editing in Toxoplasma gondii. Methods Mol Biol. 2017;1498:79–103. doi: 10.1007/978-1-4939-6472-7_6. PubMed PMID: 27709570.

106. Gibson DG, Young L, Chuang RY, Venter JC, Hutchison CA, 3rd, Smith HO. Enzymatic assembly of DNA molecules up to several hundred kilobases. Nat Methods. 2009;6(5):343–5. Epub 20090412. doi: 10.1038/nmeth.1318. PubMed PMID: 19363495.

107. Stasic AJ, Chasen NM, Dykes EJ, Vella SA, Asady B, Starai VJ, et al. The Toxoplasma Vacuolar H(+)-ATPase Regulates Intracellular pH and Impacts the Maturation of Essential Secretory Proteins. Cell Rep. 2019;27(7):2132–46 e7. doi: 10.1016/j.celrep.2019.04.038. PubMed PMID: 31091451; PubMed Central PMCID: PMCPMC6760873.

108. Liu J, Pace D, Dou Z, King TP, Guidot D, Li ZH, et al. A vacuolar-H(+) -pyrophosphatase (TgVP1) is required for microneme secretion, host cell invasion, and extracellular survival of Toxoplasma gondii. Mol Microbiol. 2014;93(4):698–712. Epub 20140716. doi: 10.1111/mmi.12685. PubMed PMID: 24975633; PubMed Central PMCID: PMCPMC4159726.

109. Schindelin J, Arganda-Carreras I, Frise E, Kaynig V, Longair M, Pietzsch T, et al. Fiji: an open-source platform for biological-image analysis. Nat Methods. 2012;9(7):676–82. Epub 20120628. doi: 10.1038/nmeth.2019. PubMed PMID: 22743772; PubMed Central PMCID: PMCPMC3855844.

110. Li ZH, Ramakrishnan S, Striepen B, Moreno SN. Toxoplasma gondii relies on both host and parasite isoprenoids and can be rendered sensitive to atorvastatin. PLoS Pathog. 2013;9(10):e1003665. Epub 20131017. doi: 10.1371/journal.ppat.1003665. PubMed PMID: 24146616; PubMed Central PMCID: PMCPMC3798403.

111. Huet D, Rajendran E, van Dooren GG, Lourido S. Identification of cryptic subunits from an apicomplexan ATP synthase. Elife. 2018;7. Epub 20180911. doi: 10.7554/eLife.38097. PubMed PMID: 30204085; PubMed Central PMCID: PMCPMC6133553.

112. Rohloff P, Miranda K, Rodrigues JC, Fang J, Galizzi M, Plattner H, et al. Calcium uptake and proton transport by acidocalcisomes of Toxoplasma gondii. PLoS One. 2011;6(4):e18390. Epub 20110425. doi: 10.1371/journal.pone.0018390. PubMed PMID: 21541023; PubMed Central PMCID: PMCPMC3081817.

113. Cho KF, Branon TC, Udeshi ND, Myers SA, Carr SA, Ting AY. Proximity labeling in mammalian cells with TurboID and split-TurboID. Nat Protoc. 2020;15(12):3971–99. Epub 20201102. doi: 10.1038/s41596-020-0399-0. PubMed PMID: 33139955.

114. Liffner B, Absalon S. Expansion Microscopy Reveals Plasmodium falciparum Blood-Stage Parasites Undergo Anaphase with A Chromatin Bridge in the Absence of Mini-Chromosome Maintenance Complex Binding Protein. Microorganisms. 2021;9(11). Epub 20211106. doi: 10.3390/microorganisms9112306. PubMed PMID: 34835432; PubMed Central PMCID: PMCPMC8620465.

115. Rinkenberger N, Rosenberg A, Radke JB, Bhushan J, Tomita T, Weiss LM, et al. Susceptibility of Toxoplasma gondii to autophagy in human cells relies on multiple interacting parasite loci. mBio. 2024;15(1):e0259523. Epub 20231214. doi: 10.1128/mbio.02595-23. PubMed PMID: 38095418; PubMed Central PMCID: PMCPMC10790690.

116. Perez-Riverol Y, Bandla C, Kundu DJ, Kamatchinathan S, Bai J, Hewapathirana S, et al. The PRIDE database at 20 years: 2025 update. Nucleic Acids Res. 2025;53(D1):D543-D53. doi: 10.1093/nar/gkae1011. PubMed PMID: 39494541; PubMed Central PMCID: PMCPMC11701690.

