## Supplemental tables and Figures for "A Protein Disulfide Isomerase Coordinates Redox Homeostasis and ER Calcium Regulation for Optimal Lytic Cycle Progression in *Toxoplasma gondii*"

Supplementary Information

**Supplementary Tables**

**Supplementary Table S1. Phylogenetic sequences used for Figs 1 and S2**

| Phylogeny | Organism | Genebank/VEuPathDB | Reference |
|---|---|---|---|
| TGGT1_211680 | *Toxoplasma gondii* | TGGT1_211680 | This work |
| | *Hammondia hammondi* | HHA_211680 | BLAST predicted |
| | *Neospora caninum* | NCLIV_011410 | BLAST predicted |
| | *Besnoitia besnoiti* | BESB_016710 | BLAST predicted |
| | *Cystospora suis* | CSUI_002988 | BLAST predicted |
| | *Sarcocystis neurona* | SN3_02200010 | BLAST predicted |
| | *Cyclospora cayetanensis* | cyc_03227 | BLAST predicted |
| | *Plasmodium knowlesi* | PKNH_1322000 | BLAST predicted |
| | *Plasmodium vivax* | PVP01_0506500 | [1] |
| | *Cryptosporidium parvum* | cgd6_120 | BLAST predicted |
| | *Saccharomyces cerevisiae* | YCL043C | [2-4] |
| | *Sporisorium reilianum* | sr10870 | BLAST predicted |
| | *Leishmania donovani* | LdBPK_367280.1 | [5] |
| | *Trypanosoma cruzi* | TcCLB.507611.370 | BLAST predicted |
| | *Trypanosoma brucei* | Tb11.v5.0668 | BLAST predicted |
| | *Gallus gallus* | Q8JG64 | [6] |
| | *Bos taurus* | P38657 | [7] |
| | *Homo sapiens* | P30101 | [8] |
| | *Mus musculus* | P27773 | [9] |
| | *Bos taurus* | Q29RV1 | PDIA4 BLAST predicted |
| | *Homo sapiens* | P13667 | [10] |
| | *Mus musculus* | P08003 | [11] |
| TGGT1_249270 | *Toxoplasma gondii* | TGGT1_249270 | This work |
| | *Hammondia hammondi* | HHA_249270 | BLAST predicted |
| | *Neospora caninum* | NCLIV_065470 | BLAST predicted |
| | *Besnoitia besnoiti* | BESB_004360 | BLAST predicted |
| | *Cystospora suis* | CSUI_008082 | BLAST predicted |
| | *Sarcocystis neurona* | SN3_00700720 | BLAST predicted |
| | *Cyclospora cayetanensis* | cyc_05674 | BLAST predicted |
| | *Plasmodium knowlesi* | PKNH_0932300 | BLAST predicted |
| | *Plasmodium vivax* | PVP01_0934900 | BLAST predicted |
| | *Cryptosporidium parvum* | cgd7_4080 | BLAST predicted |
| | *Saccharomyces cerevisiae* | YCL043C | [2-4] |
| | *Sporisorium reilianum* | sr12879 | BLAST predicted |
| | *Leishmania donovani* | LdBPK_367280.1 | [5] |
| | *Trypanosoma cruzi* | TcCLB.507611.370 | BLAST predicted |
| | *Trypanosoma brucei* | Tb11.v5.0668 | BLAST predicted |
| | *Gallus gallus* | geneid_421940 | BLAST predicted |
| | *Bos taurus* | ENSBTAG00000001928 | PDIA6 BLAST predicted |
| | *Homo sapiens* | ENSG00000143870 | [12] |
| | *Mus musculus* | ENSMUSG00000020571 | [13] |

Supplementary Table S1. All Phylogenetic sequences used, including organism, gene ID, and method for obtaining sequence. BLAST predictions were conducted on VEUpathDB.org.

**Supplementary Table S2. List of Primers used**

| Cell Line | Primer name | Sequence |
|---|---|---|
| Recombinant TgPDIA3 | 211680 gene Fwd | atgcgagccgggttttcgtttgctctgttggcagtcggcc |
| | 211680 gene Rev | ttacagttcttcacccttgtcgtccttc |
| | 211680 gene with homology Fwd | tcaccatcaccgagccgggttttcgtttg |
| | 211680 gene with homology Rev | gagtccaagcttacagttcttcacccttgtcg |
| | pQE80L with 211680 homology Fwd | agaactgtaagcttggactcctgttgatag |
| | pQE80L with 211680 homology Rev | acccggctcggtgatggtgatggtgatg |
| Recombinant TgPDIA6 | 249270 gene Fwd | atggcgttcacaggcgcg |
| | 249270 gene Rev | tcaaagttcatctttcggaagttcttcatctttg |
| | 249270 gene with homology Fwd | tcaccatcacgcgttcacaggcgcgcgc |
| | 249270 gene with homology Rev | gagtccaagctcaaagttcatctttcggaagttcttcatctttgccatcc |
| | pQE80L with 249270 homology Fwd | tgaactttgagcttggactcctgttgatag |
| | pQE80L with 249270 homology Rev | ctgtgaacgcgtgatggtgatggtgatg |
| *iΔTgPDIA3-3Ty* | 211680 PI3Ty Fwd | tcctccgtctccgtcgtacccaggaagttttccctgtcggcatatgcgtgactttcccgc |
| | 211680 PI3Ty Rev | ggccgactgccaacagagcaaacgaaaacccggctcgcatgtccaggggatcctgattg |
| | 211680 N-terminal gRNA Fwd | gctttgtctcgcttagagcctgttttagagctagaaatagcaag |
| | 211680 N-terminal validation Fwd | acacctccactgtttccggaaca |
| *iΔTgPDIA6*-3Ty | 249270 PI3Ty Fwd | tgtcgactctttcttcgctgctgacgagcacaccgcgttccatatgcgtgactttcccgc |
| | 249270 PI3Ty Rev | gaggtgcaccgcgccgcatgcgcgcgcctgtgaacgccatgtccaggggatcctgattg |
| | 249270 N-terminal gRNA Fwd | gtcccgcgcctcgaaaaattcgttttagagctagaaatagcaag |
| | 249270 N-terminal validation Fwd | cccttcctaccttctgcatttccca |
| pLIC3-HA-KDEL plasmid | pLIC-3HA-KDEL Fwd | gaactttaacccgggcatatgtagaaaagttgtaacg |
| | pLIC-3HA-KDEL Rev | atctttggcataatctggaacatcgtaaggatacg |
| TgPDIA6-3HA-KDEL | 249270-pLIC-3HA-KDEL homology Fwd | tcgcgtgcgtgggatggcaaagatgaagaacttccgattggaagtggaggacgggaattc |
| | 249270-pLIC-3HA-KDEL homology Fwd | cccatctctctgtcggaggctgtctgcaggacagcaagaattgtgttaaccggttcgact |
| *iΔTgPDIA3* (no tag) | 211680 PI no tag Fwd | ggccgactgccaacagagcaaacgaaaacccggctcgcatggttgaagacagacgaaagc |
| | 211680 PI no tag Rev | gaggtgcaccgcgccgcatgcgcgcgcctgtgaacgccatggttgaagacagacgaaagc |
| pLIC3-HA-GEEL plasmid | pLIC-3HA-GEEL Fwd | gaactttaacccgggcatatgtagaaaagttgta |
| | pLIC-3HA-GEEL Rev | ttcaccggcataatctggaacatcgtaa |
| TgPDIA3-3HA-GEEL | 6803HACFwd | aagccgctcaagaaggacgacaagggtgaagaactgattggaagtggaggacgggaattc |
| | 6803HACRev | gggttatattgacggaaacatgagccaaacagggcaagaattgtgttaaccggttcgact |

| | | |
|---|---|---|
| | 211680 C-term gRNA | ggtacatgagcggttgaaacagttttagagctagaaatagcaag |
| TurboID-3HA-GEEL plasmid | TurboID-3HA-GEEL Fwd | gaactttaacccgggcatatgtagaaaagttgtaa |
| | TurboID-3HA-GEEL Rev | ttcacccttttcggcagaccg |
| TgPDIA3-TBID-GEEL | 211680 TurboID Fwd | aagcacggttccaagccgctcaagaaggacgacaagtacccgtacgacgtcccggactac |
| | 211680 TurboID Rev | gggttatattgacggaaacatgagccaaacaggccctcgggggggcaagaattgtgttaa |
| Recombinant RFP | RFP sequence Fwd | tcaccatcacatggcgcctagggtgagc |
| | RFP sequence Rev | gagtccaagccttgtacagctcgtccatgc |
| | pQE80L with RFP homology Fwd | gctgtacaaggcttggactcctgttgatag |
| | pQE80L with RFP homology Rev | taggcgccatgtgatggtgatggtgatg |
| | RFP validation rev | ccgcgcatcttcaccttgtagatca |
| Recombinant GFP | GFP sequence Fwd | tcaccatcacatggtgagcaagggcgag |
| | GFP sequence Rev | gagtccaagcttacttgtacagctcgtccac |
| | pQE80L with GFP homology Fwd | gtacaagtaagcttggactcctgttgatag |
| | pQE80L with GFP homology Rev | tgctcaccatgtgatggtgatggtgatg |
| | GFP validation Fwd | gggcatggcggacttgaagaa |
| Recombinant GRA1 for antibody | GRA1 gene Fwd | tcaccatcacatggtgcgtgtgagcgct |
| | GRA1 gene Rev | gagtccaagcttactctctctctcctgttaggaacc |
| | pQE80L with GRA1 homology Fwd | gagagagtaagcttggactcctgttgatag |
| | pQE80L with GRA1 homology Rev | cacgcaccatgtgatggtgatggtgatg |
| | GRA1 validation Rev | tcgctgtacgatccatctgaagctttaat |
| *TgERDJ3A-3HA-KDEL* | 209950 C-3HA Fwd | aaggaagaaacagagaagaaggagaaggcagacaagattggaagtggaggacggga a |
| | 209950 C-3HA Rev | aaacgtacgacggggagagacaggaaagggaacgcaagaattgtgttaaccggttcgact |
| | 209950 C-3HA C-Val F | ttcaaggttgtgccatcgagcaaga |
| | 209950 C-Cas9 Fwd | gacatctaccggatctctgtggttttagagctagaaatagcaag |
| *iΔTgERDJ3A-3Ty* | 209950 N-Cas9 Fwd | gtgtggcggtgggggtcgctggttttagagctagaaatagcaag |
| | 209950 PI3Ty Fwd | actgtcccctcctcctcccctccccccctcctcccccgcccatatgcgtgactttcccgc |
| | 209950 PI3Ty Rev | agacgagaggcaagaacgcgcacacacgcagaggcgccatgtccaggggatcctgattg |
| | 209950 N-val Rev | gaacttctggaccttcccgtccc |
| *iΔTgERO1-3Ty* | ERO1 N-gRNA | gctcaggcgtgaccagaccaggttttagagctagaaatagcaag |
| | ERO1 PI3Ty Fwd | cattgaaaagccggcccttgggaggtcgtttcgaattcagcatatgcgtgactttcccgc |
| | ERO1 PI3Ty Rev | gacgccttctccttatttccctatactgagatccttccatgtccaggggatcctgattg |
| | ERO1 N-val Rev | aaccgaagacaacgagcgaaaatcc |
| Other validation primers | (AJ11) pLIC-3HA val Rev | ggatagccagcgtagtccggg |
| | (AJ121) pQE-80L seq Rev | cttccttagctcctgaaaatctcgcc |

**Supplementary Table S3 Hits significantly enriched in both *TgPDIA3-TID* ER and TgPDIA3-IP +DVSF fraction**

| Gene ID | Gene product | L2FC +DVSF | L2FC TID ER |
|---|---|---|---|
| TGGT1_319560 | MIC3 | 9.39 | 6.41 |
| TGGT1_201780 | MIC2 | 7.64 | 3.91 |
| TGGT1_292020 | GCC2/GCC3 | 8.37 | 4.39 |
| TGGT1_300350 | cysteine desulfurase/selenocysteine lyase family PLP dependent transferase superfamily protein | 10.23 | 4.32 |
| TGGT1_248880 | GTPase RAB7 | 2.00 | 4.32 |
| TGGT1_247350 | thioredoxin domain-containing protein | 9.76 | 3.91 |
| TGGT1_253900 | parasite porphobilinogen synthase PBGS | 8.08 | 3.91 |

Supplementary Table S3. Proteins significantly enriched in both TgPDIA3 +DVSF and TgPDIA3-TID ER fraction. Including gene ID, protein product description, and $Log_2$ ratio of the fold change for both TgPDIA3 +DVSF and TgPDIA3-TID ER fraction.

**Supplementary Figures**

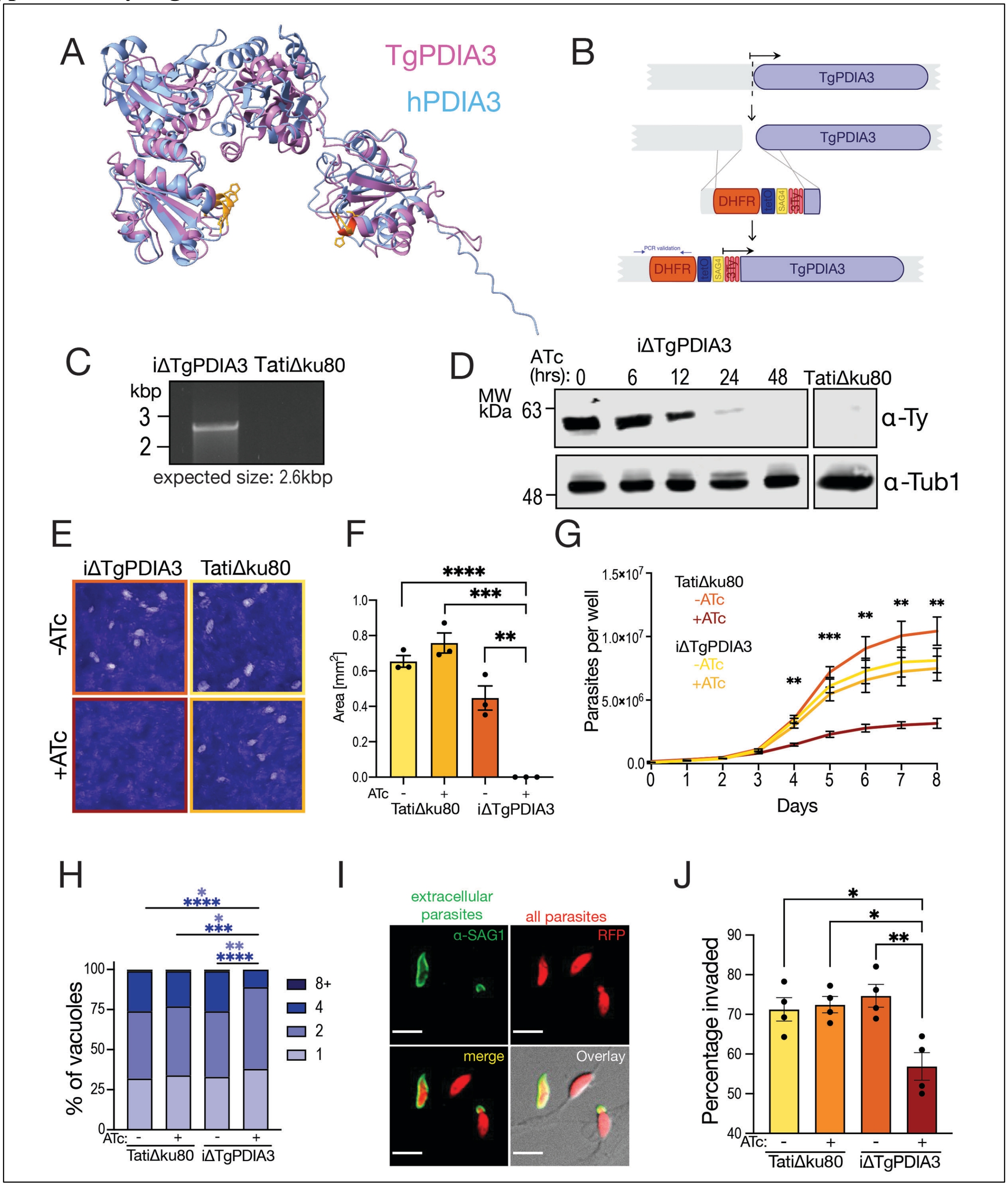


**Fig. S1. TgPDIA3 is essential for *T. gondii* growth**. A) Overlay of the TgPDIA3 and PDIA3 AlphaFold v2.0 predicted models. B) Scheme showing the promoter insertion strategy for the generation of the *iΔTgPDIA3-3Ty* mutant. DHFR, dihydrofolate reductase for pyrimethamine selection, tet operon, SAG4 promoter, and a 3xTy tag. C) PCR validation of *iΔTgPDIA3-3Ty* mutant. D) Regulation of TgPDIA3 protein expression with ATc validated by western blot. E) plaque assays of the conditional knock-down (*iΔTgPDIA3-3Ty*) mutant and *TatiΔku80* (control) incubated for 7 days with or without ATc. F) quantification of the plaque size ($mm^2$) (n=3). G) growth assay measuring parasite number standardized from RFP fluorescence of *iΔTgPDIA3-3Ty-RFP* mutant and *TatiΔku80-RFP* (control) over 8 days incubated either with or without ATc (n=3). H) quantification for replication assay, measuring the total percentage of 1, 2, 4, and 8+ parasite containing vacuoles, 20 hours post infection. I) Representative image of parasites invading host cells showing extracellular (green) and intracellular (red only) tachyzoites. J) Quantification of parasite invasion measured as the percentage of parasites invaded vs total parasites (n=4).

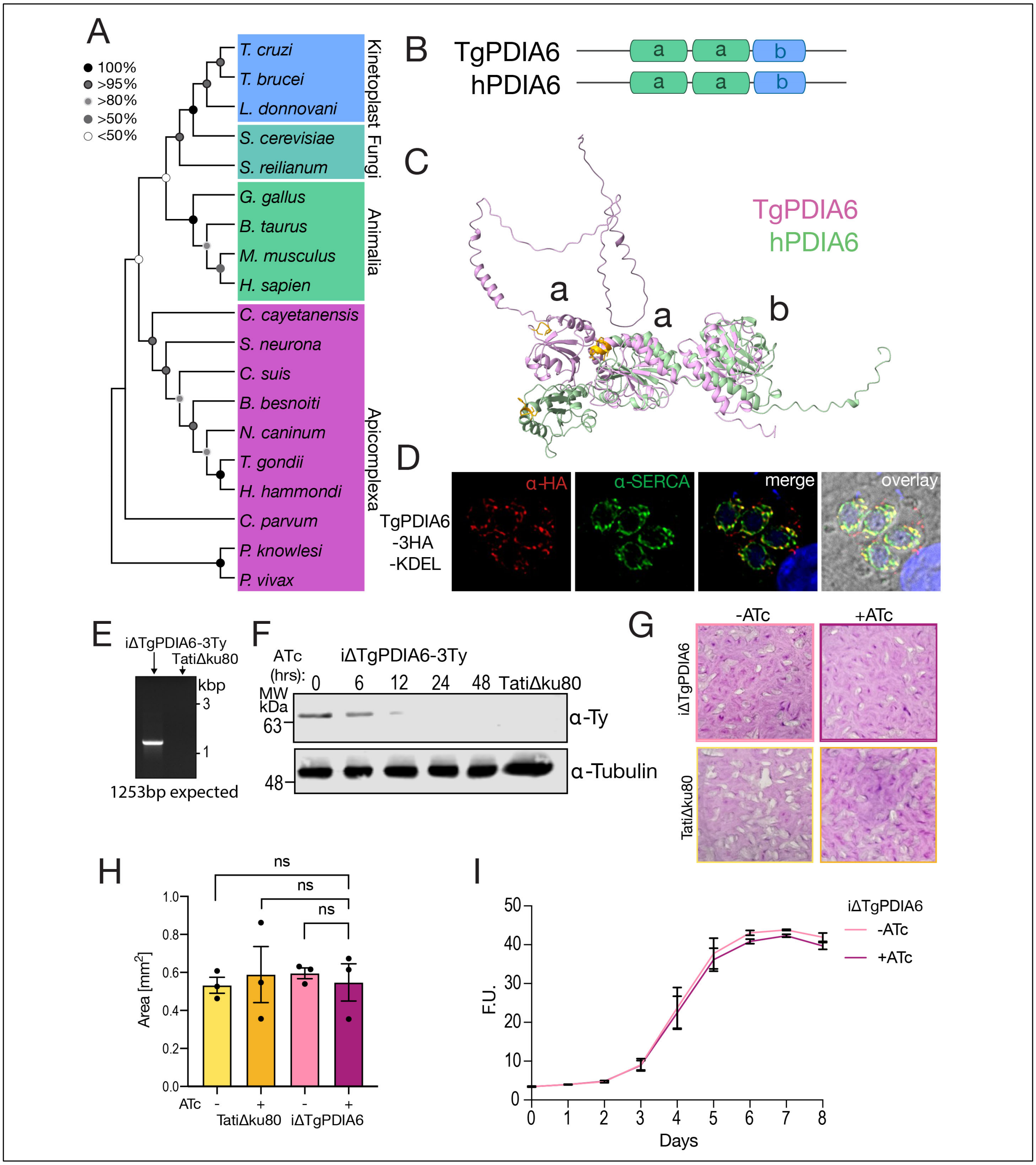


**Fig S2**. **TgPDIA6 is dispensable for *T. gondii* growth.** A) phylogenetic tree for TgPDIA6 and its blasted homologs. Sequences used are shown in Table S1. B) Catalytic a and non-catalytic b domains for TgPDIA6 and PDIA6. C) AlphaFold v2.0 model of TgPDIA6, highlighting the CXXC active site in purple. D) Immunofluorescence assay of the *TgPDIA6-3HA-KDEL* mutant probed with rat αHA (1:25) and with Guinea pig αTgSERCA (1:500). E) PCR validation for the *iΔTgPDIA6-3Ty* clone. F) Western blot showing ATc downregulation of *tgpdia6* after the addition of ATc in the *iΔTgPDIA6-3Ty* mutant. G) Representative plaque assay of *iΔTgPDIA6-3Ty* mutant and *TatiΔku80* (control) incubated for 7 days with or without ATc and H) quantification of plaque size ($mm^2$) (n=3). I) Measurement of RFP fluorescence over 8 days of the *iΔTgPDIA6-RFP* mutant incubated with or without ATc (n=3).

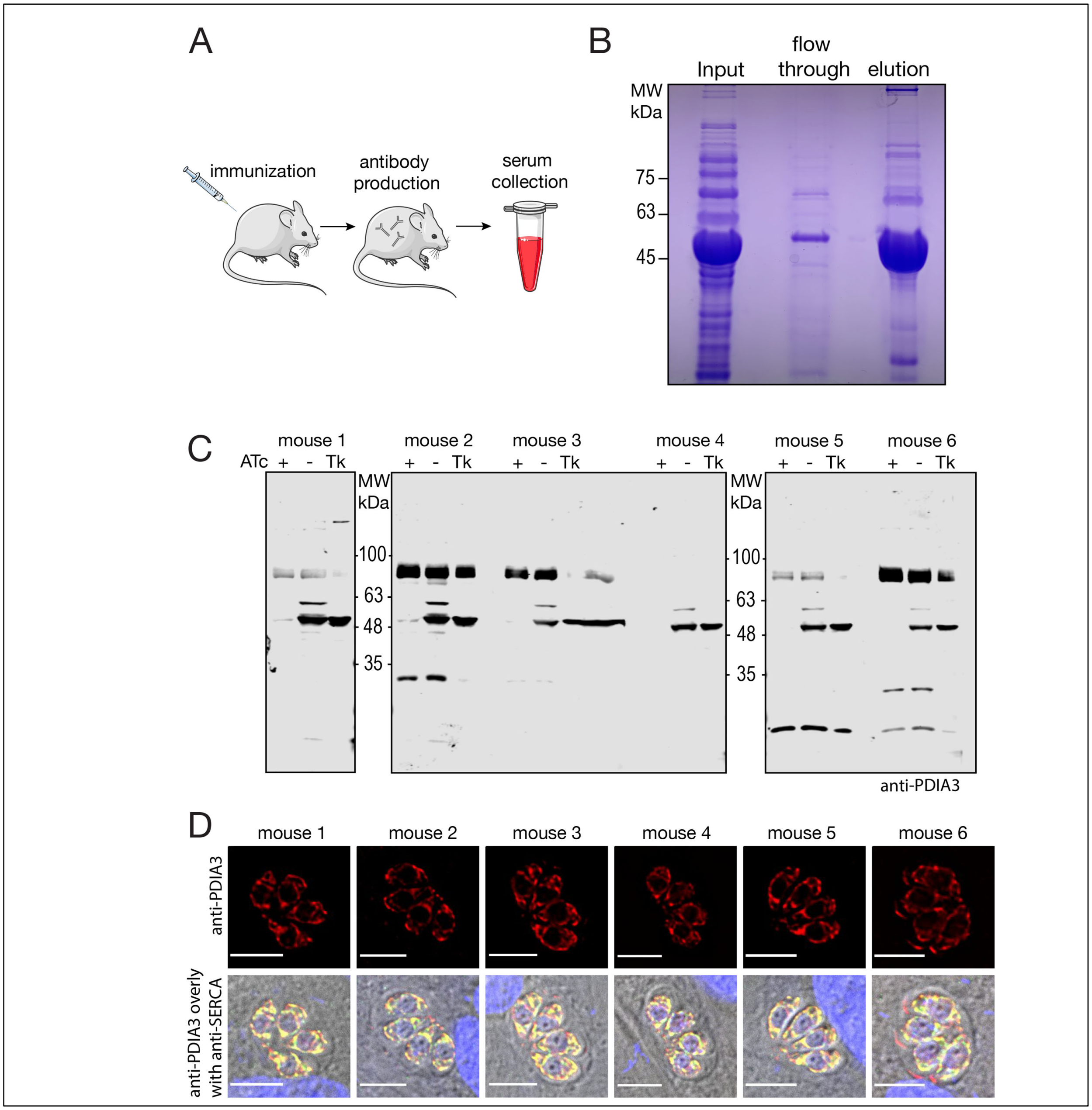


**Fig S3. Generation of the αTgPDIA3 antibody** A) cartoon depicting antibody production in mice. B) Purification of TgPDIA3 protein. C) Western blot with *TatiΔku80* (Tk) and *iΔTgPDIA3*, (-ATc and +ATc) stained with αTgPDIA3 from each of 6 mice immunized with TgPDIA3 antigen. D) Immunofluorescence assay of *TatiΔku80* parasites stained with αTgPDIA3 from each of the 6 mice immunized with TgPDIA3 antigen.

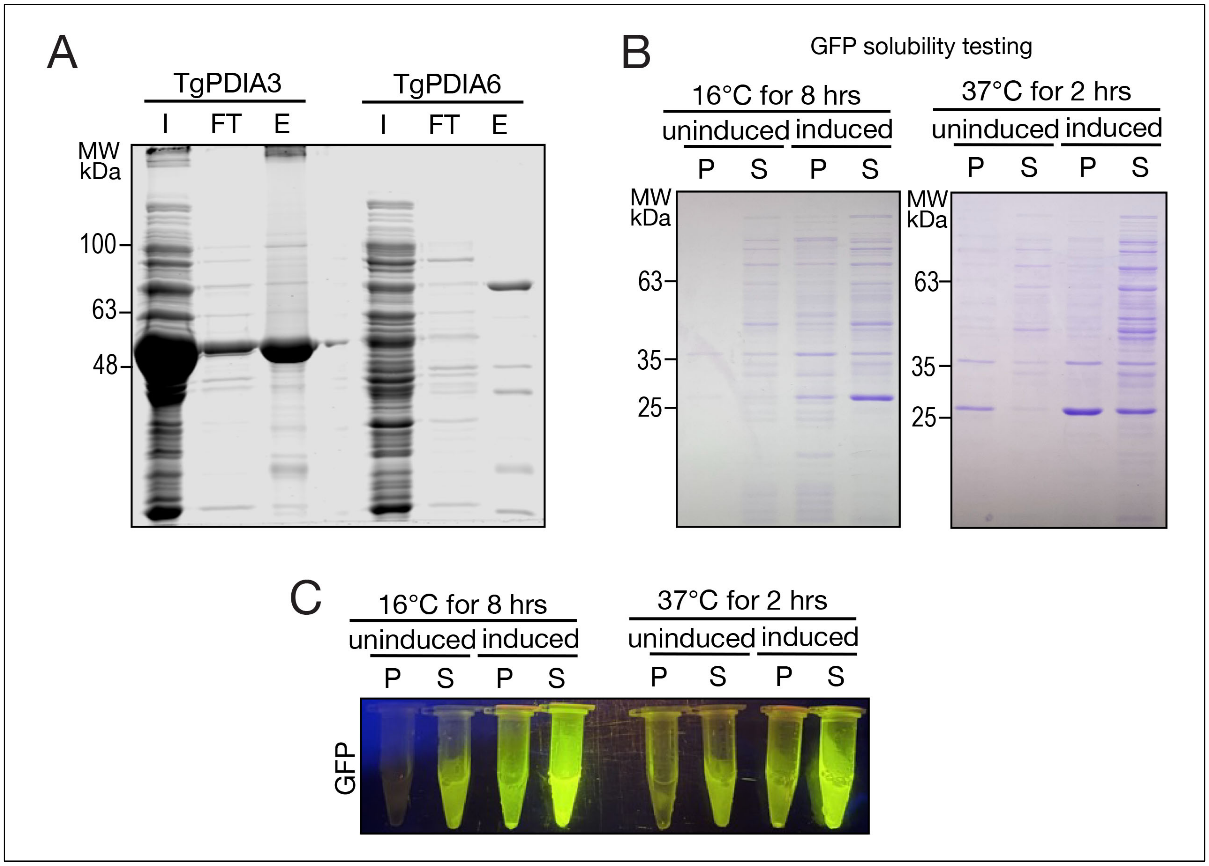


**Fig S4. Generation of recombinant GFP, TgPDIA3 and TgPDI6.** A) Coomassie-stained gel showing the protein purification of TgPDIA3 and TgPDIA6. B) Coomassie-stained gel assessing the solubility of recombinant GFP (S = soluble, P = pellet). C) Tubes of recombinant GFP in each condition showing fluorescence in the induced soluble fraction.

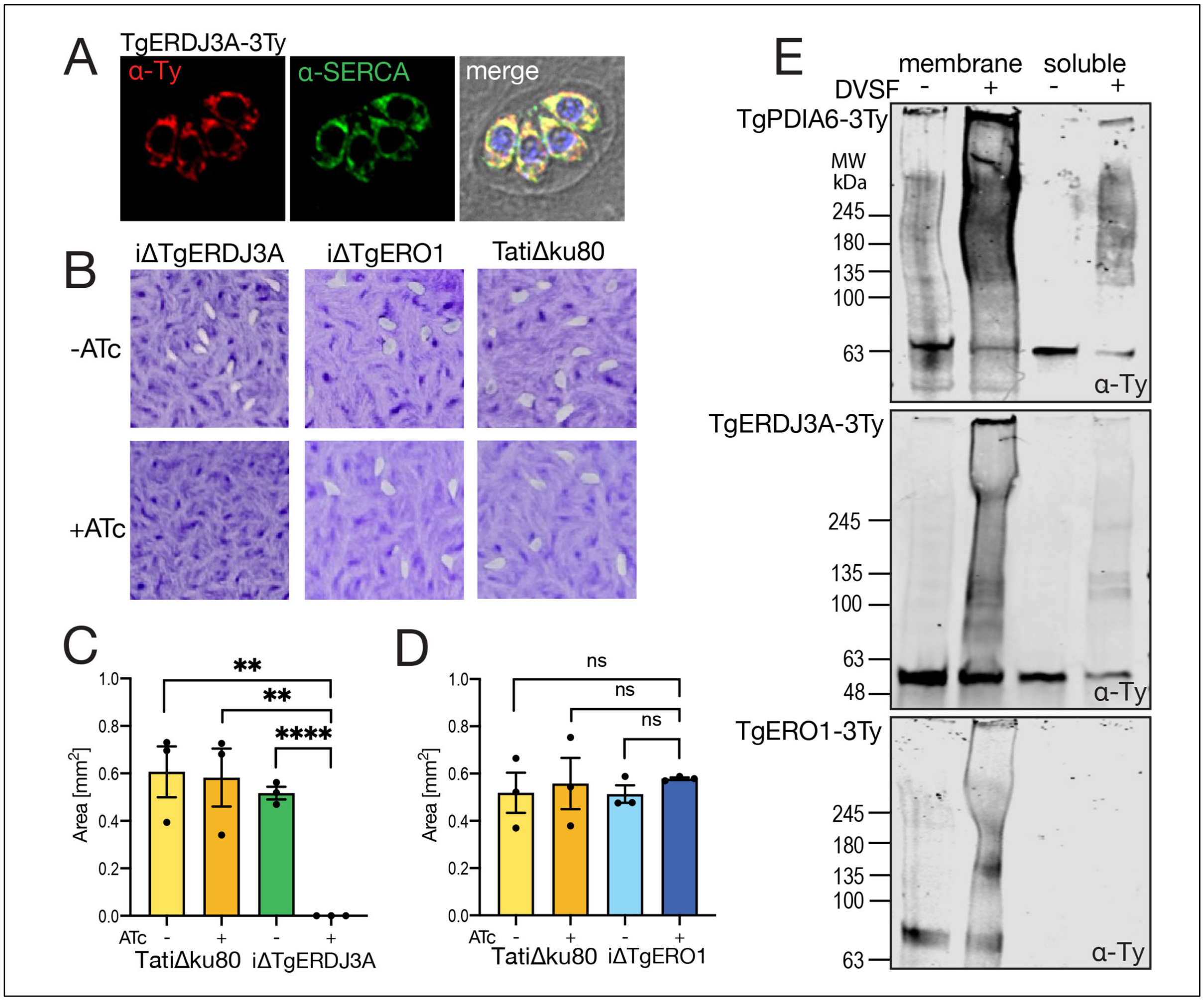


**Fig S5. TgERO1 and TgERDJ3A are ER redox enzymes.** A) Immunofluorescence assay of αTy (red) and αSERCA (green). B) representative plaque assays of *iΔTgERO1-3Ty* mutant, *iΔTgERDJ3A-3Ty* mutant and *TatiΔku80* (control) incubated for 7 days with or without ATc. Quantification of average plaque size ($mm^2$) for the *iΔTgERDJ3A-3Ty* (C) or *iΔTgERO1-3Ty* (D) mutants both including *TatiΔku80* (control) (n=3). E) Western blot of *iΔTgPDIA6-3Ty*, *iΔTgERDJ3A-3Ty*, and *iΔTgERO1-3Ty* probed with αTy, incubated either with or without DVSF for 30 min, showing either soluble proteins (right) or enriched membrane proteins (left).

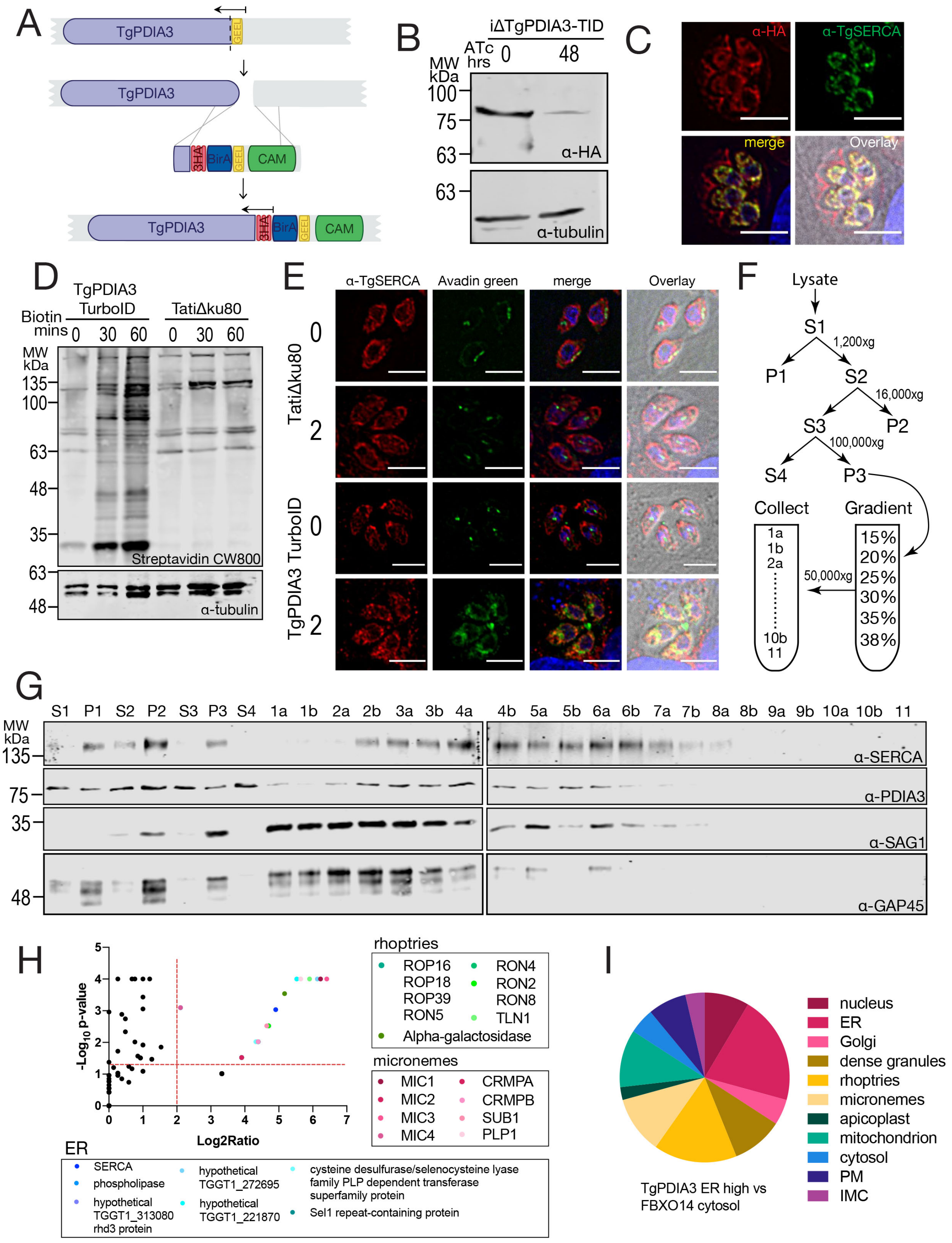


**Fig S6. Mapping the Proximal Interactome of TgPDIA3-TID.** A) Scheme showing the 3HA-TurboID cassette added to the C-terminus of *Tgpdia3,* including the GEEL retention signal downstream to the TURBO domain. A 3xHA tag is also part of the tagging. B) western blot validation of *TgPDIA3-TID-3HA-GEEL* tagging. C) IFA showing the localization of *TgPDIA3-TID-3HA-GEEL*. D) Western blot analysis of *TgPDIA3-TID-3HA-GEEL*, or parental (*TatiΔku80*) cells incubated with 50 μM biotin for 0, 30, and 60 minutes; Streptavidin shows biotinylated proteins, with tubulin as loading control. E) IFA depicting localization of biotinylated proteins in TgPDIA3-3HA-TID-GEEL compared to Tatiku80 (WT). F) scheme of the subcellular fractionation protocol. G) Western blot showing separation of all fractions depicted in (F) probed with αSERCA and αPDIA3 marking for ER proteins or αSAG1 and αGAP45 marking for plasma membrane and inner membrane complex (IMC) proteins, respectively. H) Volcano plot of proteins enriched in LC-MS/MS analysis. I) LOPIT [14]-predicted localization of proteins to the ER, microneme, or rhoptries (n=2). The complete list of peptides is in the supplemental file named Proteomic Results.

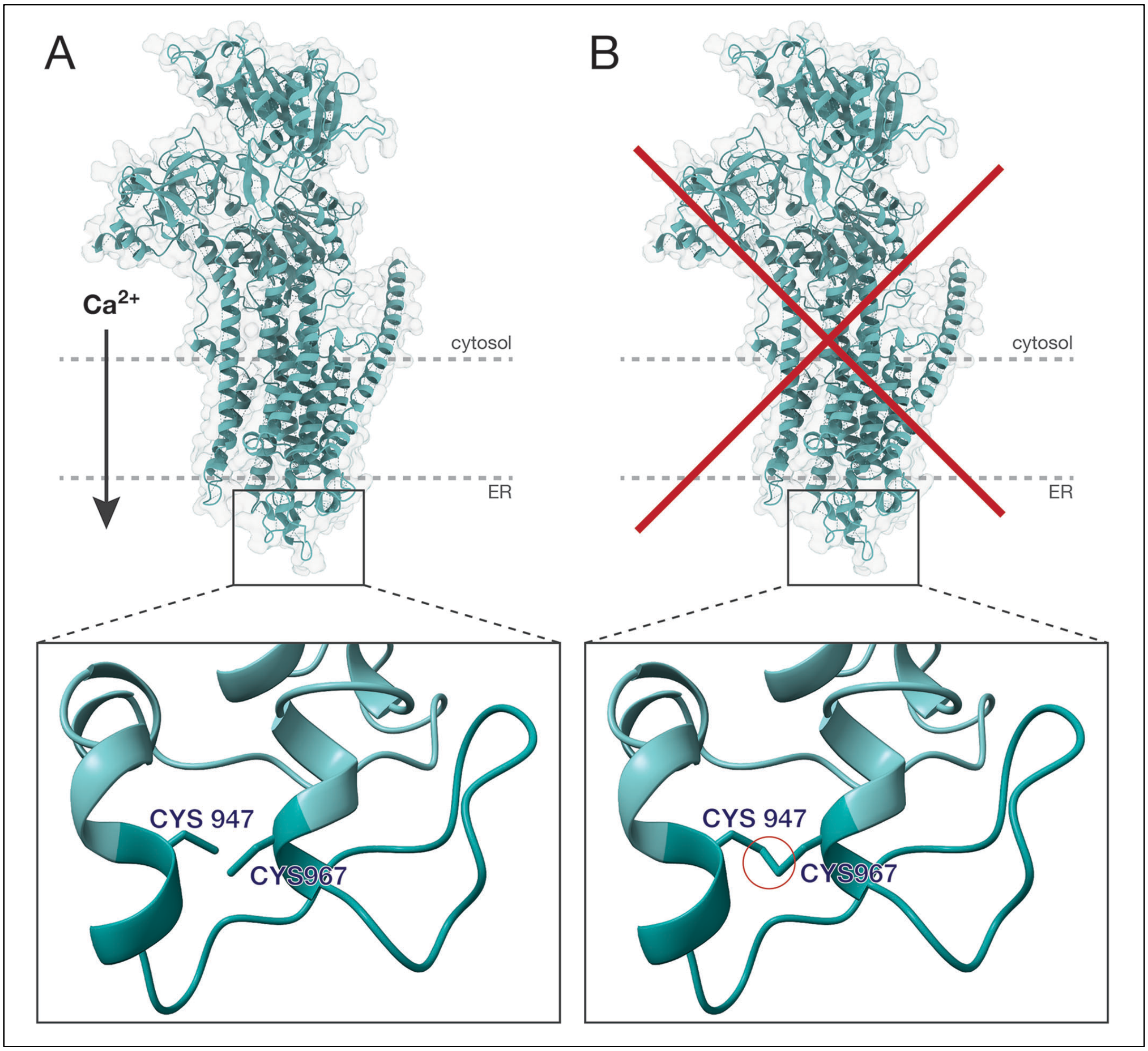


**Fig S7. Schematic of L4 luminal loop and ER redox regulation of TgSERCA.** AlphaFold v2.0 model of TgSERCA and its L4 loop, showing A) active SERCA with a reduced L4 loop B) inhibited SERCA with an oxidized L4.

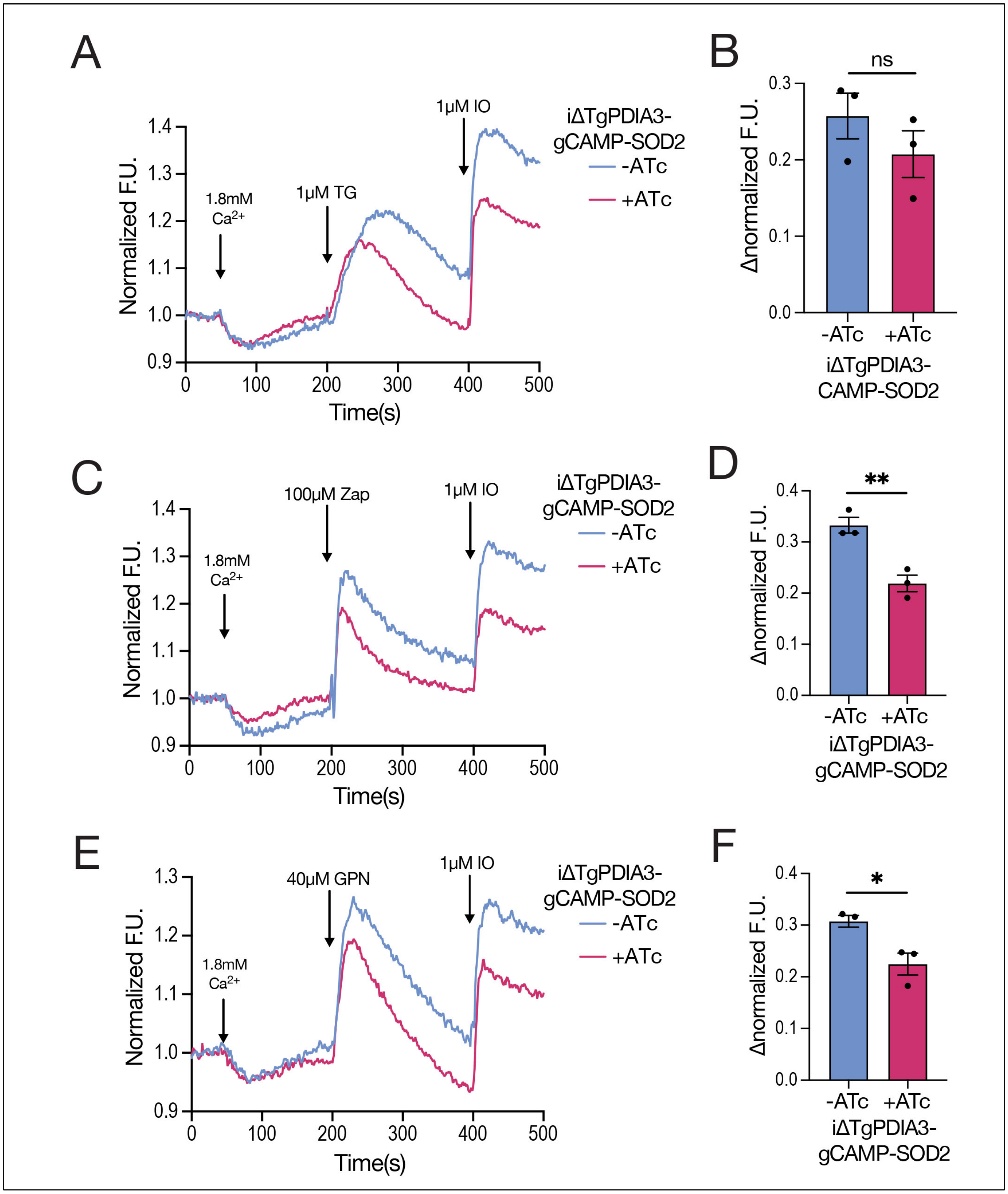


**Fig S8. Calcium uptake by the mitochondria of the *iΔTgPDIA3-GCaMP6f-SOD2* mutant.** A) Normalized mitochondrial GCaMP6 fluorescence changes in response to thapsigargin and ionomycin; B) quantification of normalized fluorescence change after addition of thapsigargin. C) Normalized mitochondrial GCaMP6 fluorescence changes in response to zaprinast and ionomycin; D) quantification of normalized fluorescence change after addition of zaprinast. E) Normalized mitochondrial GCaMP6 fluorescence changes in response to GPN and ionomycin; F) quantification of normalized fluorescence change after addition of GPN (n=3).

**Supplemental Video 1. U-ExM of *T. gondii* ER and micronemes.** Representative IMARIS v9 surfaces created from ultrastructure expansion microscopy with immunofluorescence labeling of intracellular *TatiΔku80* tachyzoites probed with αTgPDIA3 (clear red) and αMIC2 (solid blue).

### References


1. Mahajan B, Noiva R, Yadava A, Zheng H, Majam V, Mohan KV, et al. Protein disulfide isomerase assisted protein folding in malaria parasites. Int J Parasitol. 2006;36(9):1037-48. Epub 20060530. doi: 10.1016/j.ijpara.2006.04.012. PubMed PMID: 16806221.
2. Tachikawa H, Miura T, Katakura Y, Mizunaga T. Molecular structure of a yeast gene, PDI1, encoding protein disulfide isomerase that is essential for cell growth. J Biochem. 1991;110(2):306-13. doi: 10.1093/oxfordjournals.jbchem.a123576. PubMed PMID: 1761527.
3. Scherens B, Dubois E, Messenguy F. Determination of the sequence of the yeast YCL313 gene localized on chromosome III. Homology with the protein disulfide isomerase (PDI gene product) of other organisms. Yeast. 1991;7(2):185-93. doi: 10.1002/yea.320070212. PubMed PMID: 2063627.
4. LaMantia M, Miura T, Tachikawa H, Kaplan HA, Lennarz WJ, Mizunaga T. Glycosylation site binding protein and protein disulfide isomerase are identical and essential for cell viability in yeast. Proc Natl Acad Sci U S A. 1991;88(10):4453-7. doi: 10.1073/pnas.88.10.4453. PubMed PMID: 1840696; PubMed Central PMCID: PMCPMC51678.
5. Kushawaha PK, Gupta R, Tripathi CD, Sundar S, Dube A. Evaluation of Leishmania donovani protein disulfide isomerase as a potential immunogenic protein/vaccine candidate against visceral Leishmaniasis. PLoS One. 2012;7(4):e35670. Epub 20120423. doi: 10.1371/journal.pone.0035670. PubMed PMID: 22539989; PubMed Central PMCID: PMCPMC3335089.
6. Nemere I. The 1,25D3-MARRS protein: contribution to steroid stimulated calcium uptake in chicks and rats. Steroids. 2005;70(5-7):455-7. Epub 20050317. doi: 10.1016/j.steroids.2005.02.005. PubMed PMID: 15862830.
7. Hirano N, Shibasaki F, Sakai R, Tanaka T, Nishida J, Yazaki Y, et al. Molecular cloning of the human glucose-regulated protein ERp57/GRP58, a thiol-dependent reductase. Identification of its secretory form and inducible expression by the oncogenic transformation. Eur J Biochem. 1995;234(1):336-42. doi: 10.1111/j.1432-1033.1995.336_c.x. PubMed PMID: 8529662.
8. Peaper DR, Wearsch PA, Cresswell P. Tapasin and ERp57 form a stable disulfide-linked dimer within the MHC class I peptide-loading complex. EMBO J. 2005;24(20):3613-23. Epub 20050929. doi: 10.1038/sj.emboj.7600814. PubMed PMID: 16193070; PubMed Central PMCID: PMCPMC1276702.
9. Celli CM, Jaiswal AK. Role of GRP58 in mitomycin C-induced DNA cross-linking. Cancer Res. 2003;63(18):6016-25. PubMed PMID: 14522930.
10. Aguilar-Hernandez N, Meyer L, Lopez S, DuBois RM, Arias CF. Protein Disulfide Isomerase A4 Is Involved in Genome Uncoating during Human Astrovirus Cell Entry. Viruses. 2020;13(1). Epub 20201231. doi: 10.3390/v13010053. PubMed PMID: 33396308; PubMed Central PMCID: PMCPMC7824429.
11. Mazzarella RA, Srinivasan M, Haugejorden SM, Green M. ERp72, an abundant luminal endoplasmic reticulum protein, contains three copies of the active site sequences of protein disulfide isomerase. J Biol Chem. 1990;265(2):1094-101. PubMed PMID: 2295602.

12. Hayano T, Kikuchi M. Cloning and sequencing of the cDNA encoding human P5. Gene. 1995;164(2):377-8. doi: 10.1016/0378-1119(95)00474-k. PubMed PMID: 7590364.
13. Lay AJ, Dupuy A, Hagimola L, Tieng J, Larance M, Zhang Y, et al. Endoplasmic reticulum protein 5 attenuates platelet endoplasmic reticulum stress and secretion in a mouse model. Blood Adv. 2023;7(9):1650-65. doi: 10.1182/bloodadvances.2022008457. PubMed PMID: 36508284; PubMed Central PMCID: PMCPMC10182305.
14. Barylyuk K, Koreny L, Ke H, Butterworth S, Crook OM, Lassadi I, et al. A Comprehensive Subcellular Atlas of the Toxoplasma Proteome via hyperLOPIT Provides Spatial Context for Protein Functions. Cell Host Microbe. 2020;28(5):752-66 e9. Epub 20201013. doi: 10.1016/j.chom.2020.09.011. PubMed PMID: 33053376; PubMed Central PMCID: PMCPMC7670262.
